# Pragmatic analysis with knowledge-guided for unraveling peptide-protein pairwise non-covalent mechanisms

**DOI:** 10.64898/2026.02.12.705667

**Authors:** Shutao Chen, Ke Yan, Jiangyi Shao, Xiangxiang Zeng, Bin Liu

## Abstract

Understanding peptide-protein interactions is vital for decoding cellular signaling and developing targeted therapies. However, the complexity of multi-molecular associations and diverse non-covalent interactions make accurate prediction and site-specific annotation challenging. Here, we propose KGIPA, a knowledge-guided pragmatic analysis framework that incorporates pragmatic concepts from natural language into life science, capturing the influence of biological environments on non-covalent interactions. KGIPA integrates intra- and extra-linguistic contextual information to combine multimodal single-molecule features and build residue-level interaction maps. It also uses biological prior knowledge to coordinate various non-covalent interaction types. Benchmark tests demonstrate KGIPA outperforms the state-of-the-art methods in evaluating molecular binding, including protein and peptide binding residues and residue-pair interactions. Furthermore, KGIPA demonstrates strong performance in peptide-protein binding affinity prediction and peptide virtual screening. Wet-lab experiments validate its reliability, revealing high consistency between predicted and experimentally measured binding behaviors. These results highlight KGIPA’s potential to accelerate peptide drug discovery and establish pragmatic analysis as an effective paradigm for decoding the molecular language of interactions.

## 1 INTRODUCTION

Elucidating the mechanisms of peptide-protein interactions (PepPIs) is essential for uncovering cellular signaling processes and advancing the development of targeted therapeutics^1-3^. Although traditional biological experiments are reliable, identifying peptides and their binding proteins in complex conformational spaces is time-consuming and resource-intensive, limiting their applicability to large-scale datasets^4,5^. By contrast, computational methods identifying peptide-protein interactions at the residue level considerably enhance research efficiency, reducing the time and experimental workload required for drug development. Therefore, computational methods have gained increasing attention and made remarkable progress in recent years^6-9^.

Traditional structure-based computational methods, such as docking^10-12^ and molecular dynamics (MD) simulations^13,14^, have been employed to study the interactions between biomacromolecules (e.g., peptides) and proteins. These methods can predict potential binding sites and affinities, offering inherent interpretability. However, the accuracy of both docking and MD simulations depends on prior knowledge of protein structures, making them less effective when the three-dimensional (3D) structure of a protein is unknown^4^. Furthermore, docking peptides to proteins with unknown active sites or domains requires an extensive conformational search of peptide-protein complexes, often demanding substantial computational resources^15^.

Recently, the fusion of artificial intelligence (AI) and biotechnology has sparked innovations that have significantly advanced research in the life sciences^16-19^. The adaptation of deep learning techniques from natural language processing (NLP) has been a major driver of AI’s success in protein-related studies, facilitated by the striking homology between biological sequences and human language^20^. Just as sentences are composed of words arranged by syntactic rules to convey specific meanings, biological macromolecules like proteins are composed of amino acids arranged in linear chains governed by biochemical “grammars” to perform defined structural and functional roles^21^. This conceptual alignment has allowed models originally designed for textual data to be repurposed for protein modeling. For instance, AlphaFold3^22^ integrates NLP-inspired architectures to jointly model protein structures, interactions, and complexes with unprecedented accuracy. Building on foundational techniques from natural language processing, a wide range of deep learning models—including convolutional neural networks (CNNs)^7,23,24^, graph neural networks (GNNs)^6,25,26^, transformers architectures^27^, and diffusion models^22^—have been adapted to analyze complex patterns in biological sequences and have been further tailored to study peptide-protein interactions.

Despite these promising developments, existing methods based on deep learning for predicting peptide-protein interactions remain confronted with two challenges. The first challenge stems from the inherent complexity of biology, particularly in representing and integrating the multi-modal features of peptides and proteins, as well as in practically annotating their interactions. On the one hand, previous models (e.g., CAMP^7^ and IIDL-PepPI^28^) have employed a variety of features such as sequence encoding, physicochemical properties, and evolutionary information to represent peptides and proteins, but they have neglected the sequence-structure-function paradigm^29^. However, peptides or proteins with similar amino acid sequences may exhibit entirely different interaction patterns due to variations in their 3D structures^30^. On the other hand, the current methods rely on the spatial distances between peptide and protein residues as the actual label annotation for model training, which is rather crude^31^. In reality, interactions between peptides and proteins are mediated by various non-covalent bonds (e.g., hydrophobic interactions, hydrogen bonds, and water bridges), and the formation of these bonds depends on factors such as the donor and acceptor groups and the bonding angles^32^. Therefore, incomplete sequence representation can limit modeling capabilities and predictive performance. Furthermore, if the actual label annotation of non-covalent interactions is inadequate, the prediction results, even if accurate, may fail to reflect the true binding mechanisms.

The second major challenge arises from the contextual complexity inherent in peptide-protein interaction modeling. In such tasks, the functional behavior of a peptide or protein is not determined solely by its internal sequence, but is often critically influenced by the structural and biochemical context of its binding partner. Inspired by the similarities between protein sequence and natural language, our previous work introduced the concept of protein linguistic pragmatics to reveal conformational changes and functional behaviors of proteins under different biological scenarios (e.g., interactions with diverse peptides)^28^. However, our previous framework, IIDL-PepPI, considered only the linguistic context dependence while neglecting the integration of multimodal molecular features, such as predicted three-dimensional structural information. In addition, IIDL-PepPI was designed to predict binary peptide-protein interactions and identify binding residues. It did not investigate residue-residue interaction graphs or the underlying mechanisms of noncovalent interactions, including the identification of specific types of noncovalent bonds involved. To more precisely characterize the impact of context on the functions of a sequence, pragmatic analysis can be further refined into two complementary contextual layers (Figure 1)^33^. The intra-linguistic context captures dependencies among residues within a peptide or protein chain, analogous to syntactic structures in a sentence. In contrast, the extra-linguistic context reflects the regulatory influence of binding partners, conformational ensembles, and broader cellular or environmental conditions on sequence behavior, akin to discourse background or situational cues in language^34^. By explicitly modeling these two contextual layers, this comprehensive pragmatic analysis framework provides stronger support for predicting and interpreting peptide-protein interactions. Although both types of contextual information are biologically relevant, most existing models provide only a partial representation. Representative methods such as CAMP^7^, AlphaFold-Multimer^35^, and RoseTTAFold All-Atom (RFAA)^36^ capture intra-linguistic contextual dependencies or structural coevolutionary patterns, yet they do not explicitly model task-specific contextual factors necessary for interpreting interaction mechanisms. Therefore, a comprehensive and context-aware framework grounded in pragmatic analysis is needed to better represent the conditional nature of peptide-protein interactions, a direction that remains largely unexplored^37^.

**Figure 1.**
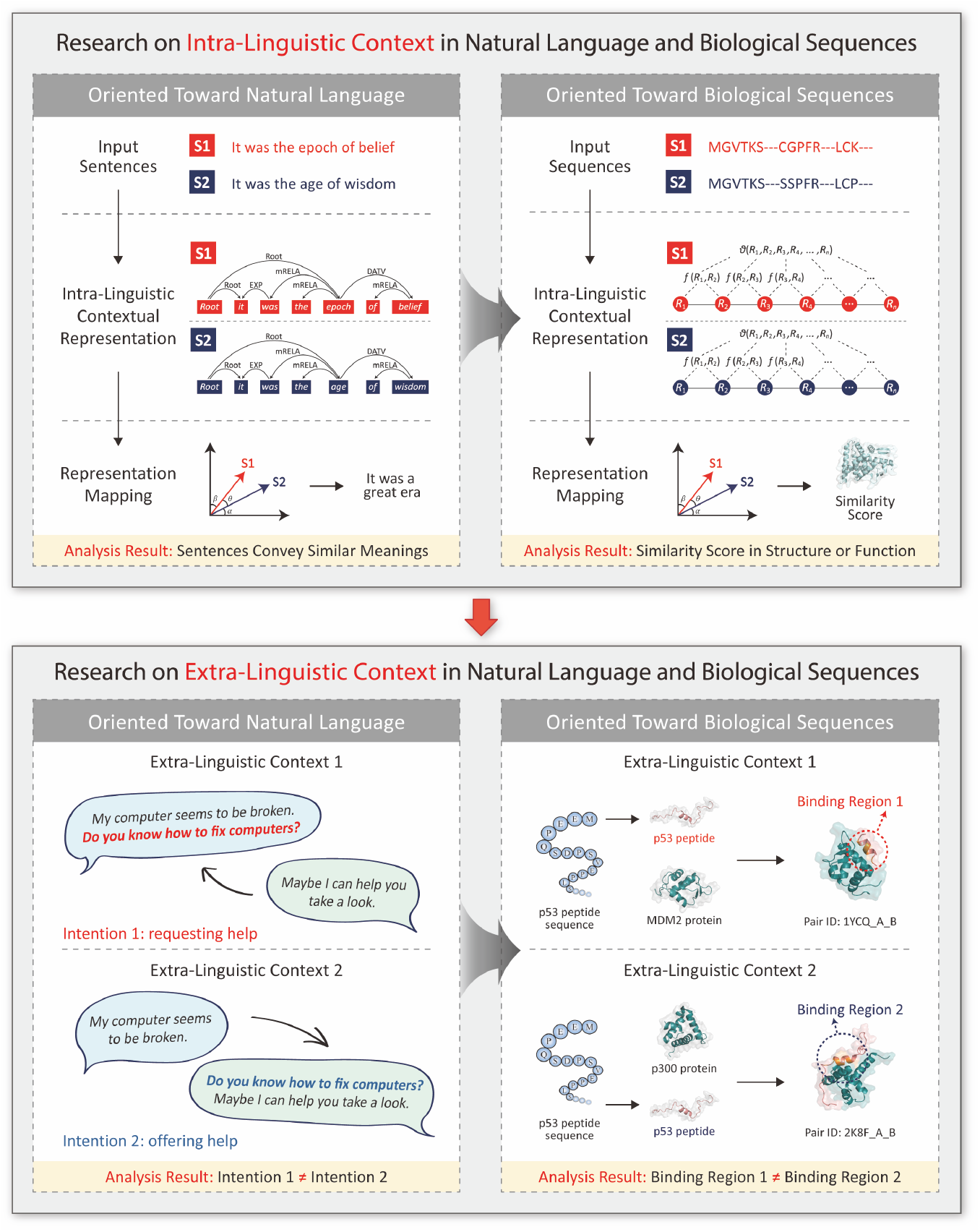
Pragmatic analysis challenges in natural language and biological sequence research: intra- and extra-linguistic contexts. Intra-linguistic context explores relationships among sequence (or sentence) monomers, while extra-linguistic context emphasizes external factors shaping sequence function (or sentence intention).

To address these challenges, we propose a knowledge-guided pragmatic analysis model (KGIPA) for predicting peptide-protein pairwise non-covalent interactions at the residue level. KGIPA is a deep learning method that integrates multi-modal features such as sequence and structure through intra-linguistic contextual representation and explicitly learns pairwise interactions via extra-linguistic contextual representation. Additionally, it employs a knowledge-guided module and a multi-task learning framework for pragmatic analysis to uncover underlying bonding mechanisms in non-covalent interactions. Compared to the state-of-the-art methods, KGIPA demonstrates superior performance and robustness in predicting peptide-protein pairwise non-covalent interactions. In peptide-protein binding affinity prediction, the pipeline from “intra-linguistic contextual representation” to “extra-linguistic contextual representation” and ultimately to “pragmatic analysis” proposed by KGIPA demonstrates strong generalizability, offering a new computational approach for AI-driven life science research. Moreover, KGIPA yields promising results in extended application tasks, including key residue identification via alanine scanning and non-covalent rule analysis.

To summarize, KGIPA distinguishes itself from previous work in three main aspects. First, it integrates multi-modal features of peptides and proteins through intra-linguistic contextual representation, comprehensively depicting individual molecules. Second, the combination of intra- and extra-linguistic contextual representations enables the fusion of peptide and protein features, effectively capturing residue-level pairwise non-covalent interactions. Lastly, KGIPA incorporates domain-specific prior knowledge through the knowledge-guided module and a multi-task learning framework, enhancing its ability to learn the underlying bonding mechanisms of non-covalent interactions for biological sequence pragmatic analysis.

## 2 RESULTS

### 2.1 Problem formulation

In peptide-protein non-covalent mechanism profiling, the task is to identify the residue pairs where a given peptide interacts with a target protein and to determine the type of non-covalent bond involved in the interaction. Given a set of peptides *M* and a set of proteins *N*, where each peptide or protein is represented by its amino acid sequence, consider the scenario where the *j*-th (*j* < |*m*|) amino acid residue of the *m*-th peptide binds with the *k*-th (*k* < |*n*|) amino acid residue of the *n*-th protein, forming the *t*-th non-covalent bond. The matrix **Y**^*t*^ can represent this relationship, where 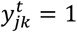denotes the presence of the bond. Since there may be *T* types of non-covalent bonds between the peptide and protein, the overall non-covalent interactions, without distinguishing between bond types, can be expressed as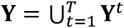. Therefore, the goal of peptide-protein non-covalent mechanism profiling is to learn a model that maps peptide and protein features to an association matrix for each type of non-covalent bond, and then combines these matrices to generate a comprehensive non-covalent interaction matrix **Y**. In this matrix **Y**, each element represents the probability score *p* of a non-covalent interaction between residue pairs, where *p* ∈ [0,1].

### 2.2 KGIPA architecture

The proposed KGIPA architecture is illustrated in Figure 2. KGIPA is designed explicitly for biological sequence pragmatic analysis, consisting of two main parts: intra-linguistic contextual representation, which focuses on individual molecules, and extra-linguistic contextual representation, which emphasizes interactions between multiple molecules. Specifically, for a given peptide-protein pair, feature extraction is first performed using various bioinformatics tools (e.g., PSI-BLAST^38^, IUPred2A^39^, SCRATCH^40^, ProtT5^17^ and trRosetta^41,42^, etc.). Next, GCN-based intra-linguistic contextual encoders integrate multi-modal features, including sequence and structure, and map them to generate intra-linguistic contextual representations corresponding to different non-covalent bonds. Subsequently, extra-linguistic contextual encoders that employ bilinear interaction mechanisms merge the peptide and protein representations, allowing the model to learn residue-level pairwise interactions. To improve the model’s ability to capture relationships between peptide-protein non-covalent bonds, we embed a knowledge-guided module, refining pairwise representations for different non-covalent mechanisms. Finally, prediction scores indicating the probability of various non-covalent interactions are generated through a series of fully connected layers, uncovering implicit non-covalent mechanisms and thereby enabling biologic sequence pragmatic analysis.

**Figure 2.**
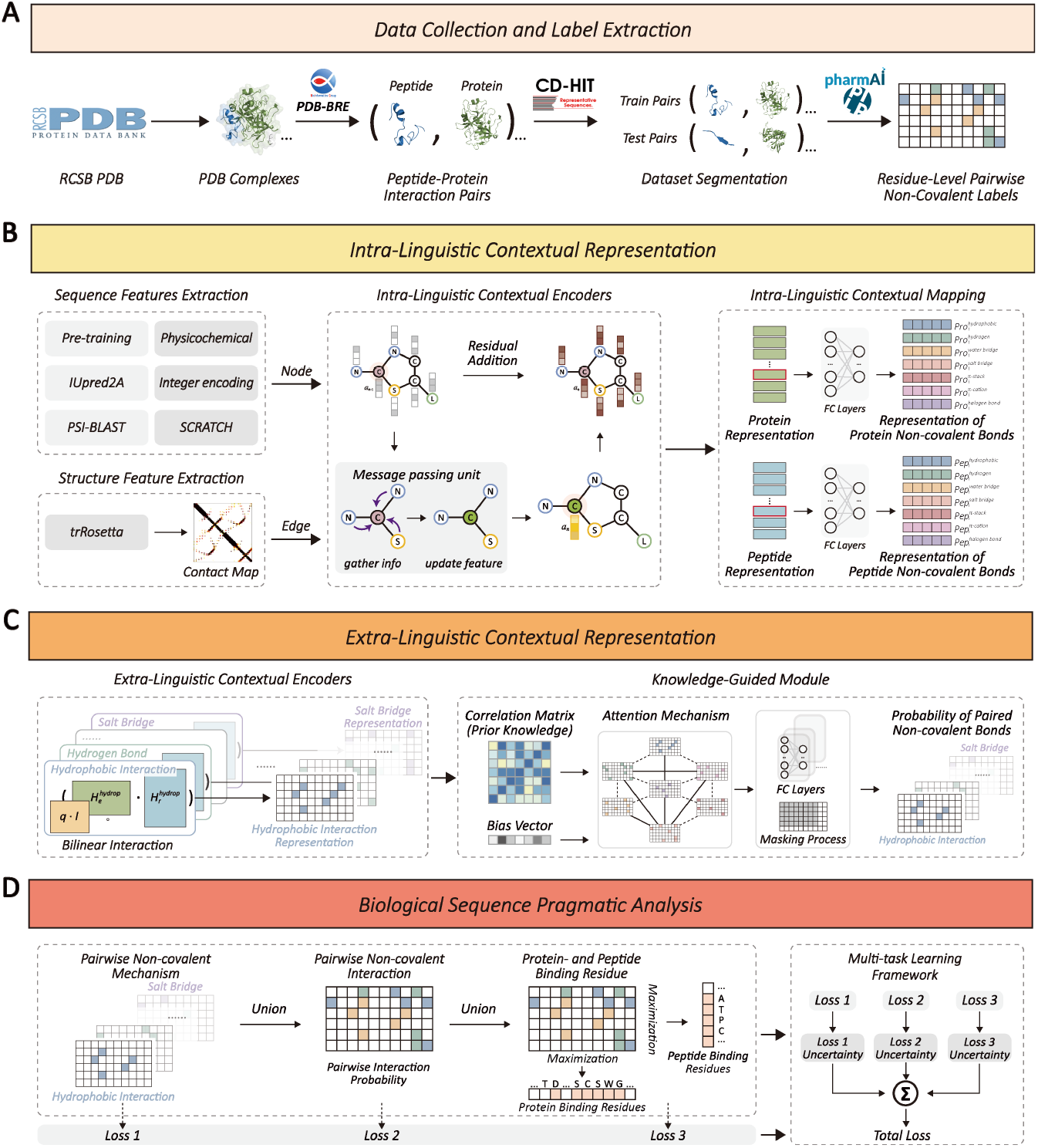
KGIPA Model Architecture. (A) Data collection and label extraction. Peptide-protein pairs were curated from the RCSB PDB using PDB-BRE, with CD-HIT ensuring sequence similarity between train and test pairs, and non-covalent bond labels derived using PLIP. (B) Intra-linguistic contextual representation. Peptide and protein multimodal features were integrated with GCN, generating representations specific to each type of non-covalent bond. (C) Extra-linguistic contextual representation. Bilinear interaction networks combined peptide and protein representations to model residue-level pairwise interactions, integrating prior knowledge of non-covalent bond types. (D) Biological sequence analysis. Multi-task learning enabled profiling of biological processes, encompassing pairwise non-covalent mechanisms, interactions, and binding residues.

### 2.3 Experimental setting

Although numerous peptide-protein interaction databases exist, most rely on spatial distance thresholds (e.g., a 5Å cutoff) for coarse interaction identification (e.g., PepBDB^31^) or do not provide residue-level binding pair information (e.g., Propedia v2.3^43^). To address this, we meticulously collected and organized peptide-protein complex data released before July 2024 based on the RCSB PDB database^44^. Peptide-protein interactions were initially identified through PDB-BRE^45^, and following Lei *et al*.^7^, training pairs with high sequence similarity (≥ 80%) to the independent test datasets were removed using CD-HIT^46,47^. The 80% sequence identity threshold is widely adopted in protein interaction studies to reduce redundancy while retaining sufficient training data, facilitating fair comparison with related methods ^48-50^. Then, the non-covalent bond type labels were extracted with PLIP^32^. Our evaluation strategy focused on independent test datasets. Specifically, we merged two previously released independent test datasets, Test251^51,52^ and LEADS-PEP^53^, and utilized the publicly available independent test dataset Test167 (comprising complexes published from September to December 2022). Additionally, we constructed a new incremental independent test dataset, Test1440, which includes complexes published from January 2023 to July 2024, to evaluate generalization performance. For further assessment of generalization under lower sequence similarity, we also derived subsets from the unified test set using more stringent CD-HIT thresholds of 60% and 40%. Notably, 40% represents the lowest sequence identity threshold supported by CD-HIT^47^. We evaluate the model across several dimensions for each independent test dataset, including peptide- and protein-binding residues, pairwise non-covalent interactions, and peptide- and protein-side non-covalent bond types. For more details on the datasets, please refer to Table S1.

For the prediction of peptide- and protein-binding residues and pairwise non-covalent interactions, we employed several commonly used evaluation metrics for binary classification tasks^54^, including Matthew’s Correlation Coefficient (MCC), Accuracy (ACC), F1 Score, Area Under the Receiver Operating Characteristic Curve (AUROC), and the Area Under the Precision-Recall Curve (AUPR). Due to the inherent class imbalance in collected dataset, AUPR was chosen as the primary evaluation metric, as it is more sensitive to the identification of positive samples and provides a clearer assessment of model performance when positive residues are relatively rare^55^. In contrast, metrics such as ACC or AUROC can be less informative under severe class imbalance, as they are often dominated by the majority class^56^. In evaluating KGIPA’s ability to predict non-covalent bond types, we used evaluation metrics for multi-class classification^57^, including Average Precision (AP), Micro F1, Micro AUROC, and Hamming Loss (HL). Five-fold cross-validation was carried out on the training dataset with a consistent random seed, selecting the model that performed best on the validation dataset. Final performance metrics were obtained by averaging the prediction results across these cross-validated models on independent test datasets.

### 2.4 Performance comparison of protein- and peptide-binding residue identification and pairwise binding prediction

Here, we merged the labels of different non-covalent bond types to directly evaluate the model’s ability to predict overall non-covalent interactions. Specifically, if any type of non-covalent bond exists between a peptide-protein residue pair, the pair is classified as having a non-covalent interaction, and the corresponding peptide and protein residues are designated as non-covalent interaction residues (also referred to as binding residues). We compared KGIPA with several SOTA baselines—RFAA^36^, PepNN^51^, CAMP^7^, MONN^25^, IIDL-PepPI^28^, and AlphaFold3^22^—using the independent test datasets Test251&LEADS-PEP, Test167, and Test1440. The five-fold cross-validation results for CAMP, MONN, IIDL-PepPI, and KGIPA are detailed in Table S2.

For the prediction of protein- and peptide-binding residues, KGIPA achieved excellent performance on both independent test datasets, Test251&LEADS-PEP (Figure 3A) and Test167 (Figure 3B). It is worth noting that PepNN only supports the prediction of protein-binding residues and therefore cannot be evaluated for peptide-binding residues, whereas CAMP has the opposite limitation. Moreover, since the training dataset of the AlphaFold3 server includes information from the Test251&LEADS-PEP dataset, we provide comparison results based on this dataset for reference. However, a rigorous evaluation of AlphaFold3 should be conducted using Test167 and Test1440.

**Figure 3.**
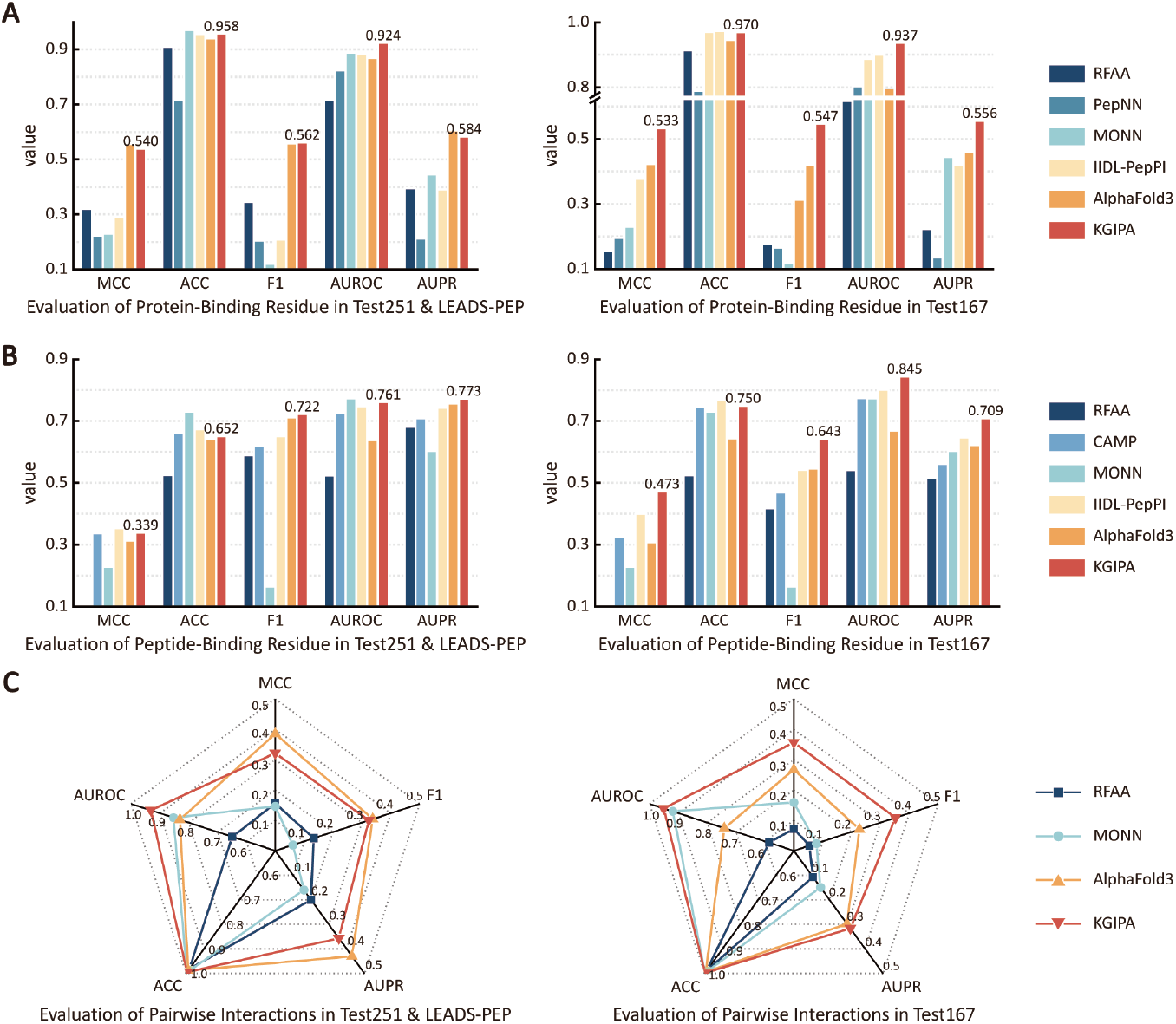
Generalization performance evaluation of KGIPA without distinguishing non-covalent interaction types. (A) Assessment of protein-binding residue identification using the independent test datasets Test251&LEADS-PEP and Test167. (B) Assessment of peptide-binding residue identification using the independent test datasets Test251&LEADS-PEP and Test167. (C) Residue-level pairwise prediction evaluation based on Test251&LEADS-PEP and Test167.

On Test167, we further conducted a one-sample t-test^58^ by comparing the AUPR values obtained from five-fold cross-validation of KGIPA with the single reported AUPR of AlphaFold3 (H0: AUPR of KGIPA ≤ AUPR of AlphaFold3). The results showed significant improvements of KGIPA over AlphaFold3 in both tasks: protein-binding residue recognition (*p*-value = 1.21×10^-5^) and peptide-binding residue recognition (*p*-value = 0.005), both rejecting the null hypothesis H0. These statistical results further confirm the advantage of KGIPA over AlphaFold3. Unlike AlphaFold3, which primarily relies on traditional techniques such as multiple sequence alignment (MSA) and template matching^59^, KGIPA benefits from a more comprehensive contextual representation framework. Furthermore, KGIPA’s superior performance can be attributed to its focus on refining peptide-protein interactions down to the residue-level non-covalent mechanisms rather than merely considering spatial physical distances. We further compared the performance of RFAA, MONN, AlphaFold3, and KGIPA in predicting peptide-protein pairwise non-covalent interactions at the residue level. As shown in Figure 3C, KGIPA achieved comparable performance to AlphaFold3 (for Test167 *p*-value = 0.460) while clearly outperforming MONN (for Test167 *p*-value = 2.07×10^-7^, for Test1440 *p*-value = 3.38×10^-6^) in residue-level pairwise interaction prediction. A comparison of AUPR values indicates that predicting pairwise interactions is more challenging than predicting peptide- and protein-binding residues. This increased difficulty arises from the more imbalanced distribution of positive and negative samples in pairwise interaction predictions, which also leads to an ACC score near 1 for different methods in residue-level pairwise interaction prediction (where the prediction tends to favor the majority negative class).

Considering recent advances in protein language models, we compared KGIPA with TAPE^60^, ProtBert^17^, ProtT5^17^, ESM 2^61^, and the latest ESM 3^62^ for protein- and peptide-binding residue prediction on Test167. As shown in Table S3, results from the independent test dataset Test167 demonstrate that KGIPA outperforms current protein language models in identifying binding residues for specific peptide-protein pairs. This superiority is attributed to KGIPA’s ability to effectively integrate contextual information through extra-linguistic contextual representation, enabling it to focus on binding residues within specific peptide-protein pairs rather than on globally significant binding residues within individual peptides or proteins. Moreover, as shown in Table S3, the protein language model ProtT5 excels at identifying the globally significant binding residues of proteins and peptides. KGIPA, building on this foundation, improves the intra-linguistic contextual representation by utilizing the pre-trained embeddings from ProtT5, thereby improving the characterization of proteins and peptides. The comparison between KGIPA and protein language models based on Test251&LEADS-PEP and Test1440 is detailed in Tables S4 and S5. For Test1440, a one-sample t-test was performed comparing the AUPR values from five-fold cross-validation of KGIPA with the single reported AUPR of AlphaFold3. KGIPA significantly outperformed AlphaFold3 in both tasks: protein-binding residue recognition (*p*-value = 0.005) and peptide-binding residue recognition (*p*-value = 7.94×10^-5^).

In addition, we clustered and partitioned the datasets using Smith-Waterman (SW) local alignment^63^ to evaluate the generalization ability of KGIPA and existing methods under three scenarios: novel peptides, novel proteins, and novel peptide-protein pairs (Note S1). The results show that as the clustering criteria become more stringent, the generalization performance of all methods decreases (Figure S1). Nevertheless, KGIPA consistently remains superior across all three scenarios, indicating its ability to effectively capture peptide-protein interaction patterns and to generalize well to unseen data.

### 2.5 Performance comparison with distinguishing non-covalent bonds

The key distinction between KGIPA and other peptide-protein interaction prediction methods lie in its ability to identify whether a specific peptide-protein residue pair forms a certain type of non-covalent interaction during binding (e.g., hydrogen bond, hydrophobic interaction). This pairwise and interaction-type-specific formulation allows KGIPA to capture fine-grained interaction patterns between peptides and proteins. We compared KGIPA with other general multi-class classification methods^64-66^ to evaluate this capability, as shown in Table 1. The results indicate that KGIPA surpasses other general models, delivering more robust predictions. Specifically, on the independent dataset Test167, KGIPA achieved lower Hamming loss for identifying non-covalent bond types on the protein side (0.083) and the peptide side (0.099) compared to other methods.

**Table 1.** Comparison between KGIPA and other SOTA multi-label learning methods.

| Methods | Evaluation Metrics |  |  |  |
| --- | --- | --- | --- | --- |
| | AP ↑ | Micro F1 ↑ | Micro AUROC ↑ | HL ↓ |
| Protein Non-Covalent Bond Type Identification |  |  |  |  |
| ML-KNN | 0.868±0.003 | 0.723±0.004 | 0.951±0.002 | 0.090±0.001 |
| Rank-SVM | 0.847±0.003 | 0.683±0.003 | 0.935±0.002 | 0.102±0.001 |
| HSCNN | 0.866±0.005 | 0.720±0.010 | 0.951±0.003 | 0.091±0.003 |
| KGIPA | **0.930±0.001** | **0.770±0.003** | **0.962±0.001** | **0.083±0.001** |
| Peptide Non-Covalent Bond Type Identification |  |  |  |  |
| ML-KNN | 0.876±0.003 | 0.721±0.006 | 0.938±0.001 | 0.116±0.002 |
| Rank-SVM | 0.876±0.004 | 0.689±0.006 | 0.926±0.002 | 0.123±0.002 |
| HSCNN | 0.874±0.003 | 0.719±0.003 | 0.937±0.002 | 0.117±0.002 |
| KGIPA | **0.910±0.003** | **0.768±0.004** | **0.953±0.001** | **0.099±0.002** |

The results are presented as mean ± standard deviation. The best result for each metric is marked in bold.

### 2.6 Effect of class imbalance and sequence similarity on prediction performance

Peptide-protein binding residue prediction is inherently affected by class imbalance, as only a small fraction of residues participates in binding interactions. To systematically assess the robustness of the proposed method under different class imbalance conditions, we conducted additional evaluations using test datasets with varying positive-to-negative sample ratios. Specifically, four independent test datasets, Test251, LEADS-PEP, Test167, and Test1440, were merged to construct a unified test dataset comprising 1,988 peptide-protein pairs. Given the substantial disparity between binding and non-binding residues, random under-sampling of the negative class was applied during evaluation to generate test sets with different positive-to-negative ratios. All competing methods were evaluated under identical sampling strategies and evaluation settings to ensure a fair comparison. As shown in Figure 4, KGIPA consistently outperforms the state-of-the-art methods across all class imbalance settings in peptide-binding residue prediction, protein-binding residue prediction, and pairwise residue interaction prediction tasks. Under the balanced condition with a positive-to-negative ratio of 1:1, KGIPA achieves improvements in AUROC of 15.2%, 14.9%, and 20.8% over AlphaFold3 for peptide, protein, and pairwise binding residue prediction, respectively. As the proportion of negative samples decreases, the ACC of all methods declines. In contrast, AUROC remains stable, and

**Figure 4.**
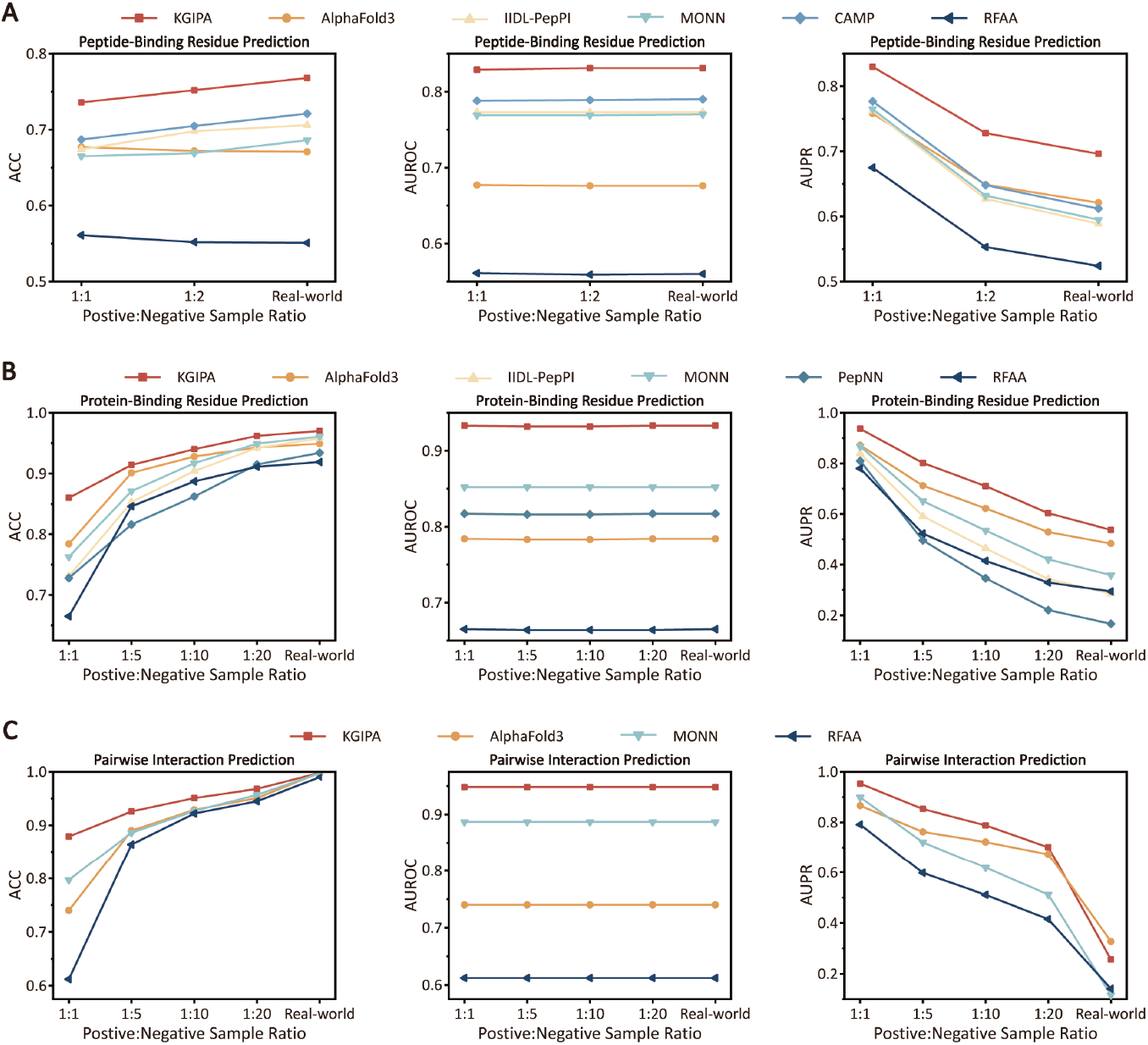
Performance of KGIPA and state-of-the-art methods under different positive-to-negative ratios. (A) Peptide-binding residue prediction under different positive-to-negative ratios. (B) Protein-binding residue prediction under different positive-to-negative ratios. (C) Pairwise interaction prediction under different positive-to-negative ratios.

AUPR increases. This phenomenon arises from changes in class balance and may affect the interpretation of certain evaluation metrics, particularly ACC, which is sensitive to class distribution^67^. Specifically, ACC is sensitive to class proportions and becomes more stringent when majority-class dominance is removed. In contrast, AUROC reflects the ranking ability of the model and is largely insensitive to class imbalance, explaining its stability. AUPR, which focuses on positive-class performance, benefits from a balanced class distribution, leading to higher values on the balanced dataset^68,69^. Nevertheless, KGIPA consistently outperforms or remains comparable to competing methods across all three metrics, demonstrating robust performance under varying class distributions. The balanced test dataset represents a theoretical scenario for evaluating the model’s intrinsic discriminative ability, whereas the original imbalanced test dataset reflects real-world data distributions. Together, these two evaluation settings provide complementary perspectives on model performance. Across both scenarios, KGIPA consistently demonstrates robust performance, outperforming competing methods.

To assess the impact of sequence similarity on prediction performance, we applied three sequence identity thresholds (80%, 60%, and 40%) between the training and test datasets by using CD-HIT to generate subsets from this unified test dataset. Please note that 40% is the lowest sequence identity threshold supported by CD-HIT^47^. To avoid effects introduced by class imbalance, all evaluations were conducted on balanced datasets. The results (Figure S2) show that as the sequence similarity threshold becomes more stringent, the performance of all methods generally declines, which is expected due to the increased difficulty of generalizing to less similar and previously unseen molecules^70^. Nevertheless, KGIPA consistently outperforms the state-of-the-art methods across all thresholds. Specifically, under the 80% threshold, KGIPA achieves AUROC improvements of 15.2%, 14.9%, and 20.8% over AlphaFold3 for peptide-binding residues, protein-binding residues, and pairwise residue interactions, respectively. At 60%, the corresponding improvements are 14.5%, 24.2%, and 33.4%, and at 40%, they remain substantial at 11.6%, 21.2%, and 29.7%. Overall, these results demonstrate that KGIPA exhibits superior generalization ability compared with existing methods.

### 2.7 Analysis of KGIPA’s knowledge-guided pragmatic framework and module contributions

The design of KGIPA follows a hierarchical modeling workflow, progressing from intra-molecular feature integration to cross-molecular representation, providing a clear rationale for how different types of information are incorporated. Specifically, peptide and protein features are first encoded independently using the intra-linguistic contextual representation, and then combined through the extra-linguistic contextual representation to model interactions between molecules. This staged, modular approach enables a transparent understanding of how each component contributes to prediction performance. We visualized intermediate features using t-SNE^71^ and observed that peptide features of binding and non-binding residues became more separable after intra-linguistic encoding, with the separation further enhanced after incorporating extra-linguistic information (Figure 5A). These results indicate that KGIPA captures biological patterns associated with known binding sites while enabling accurate classification.

**Figure 5.**
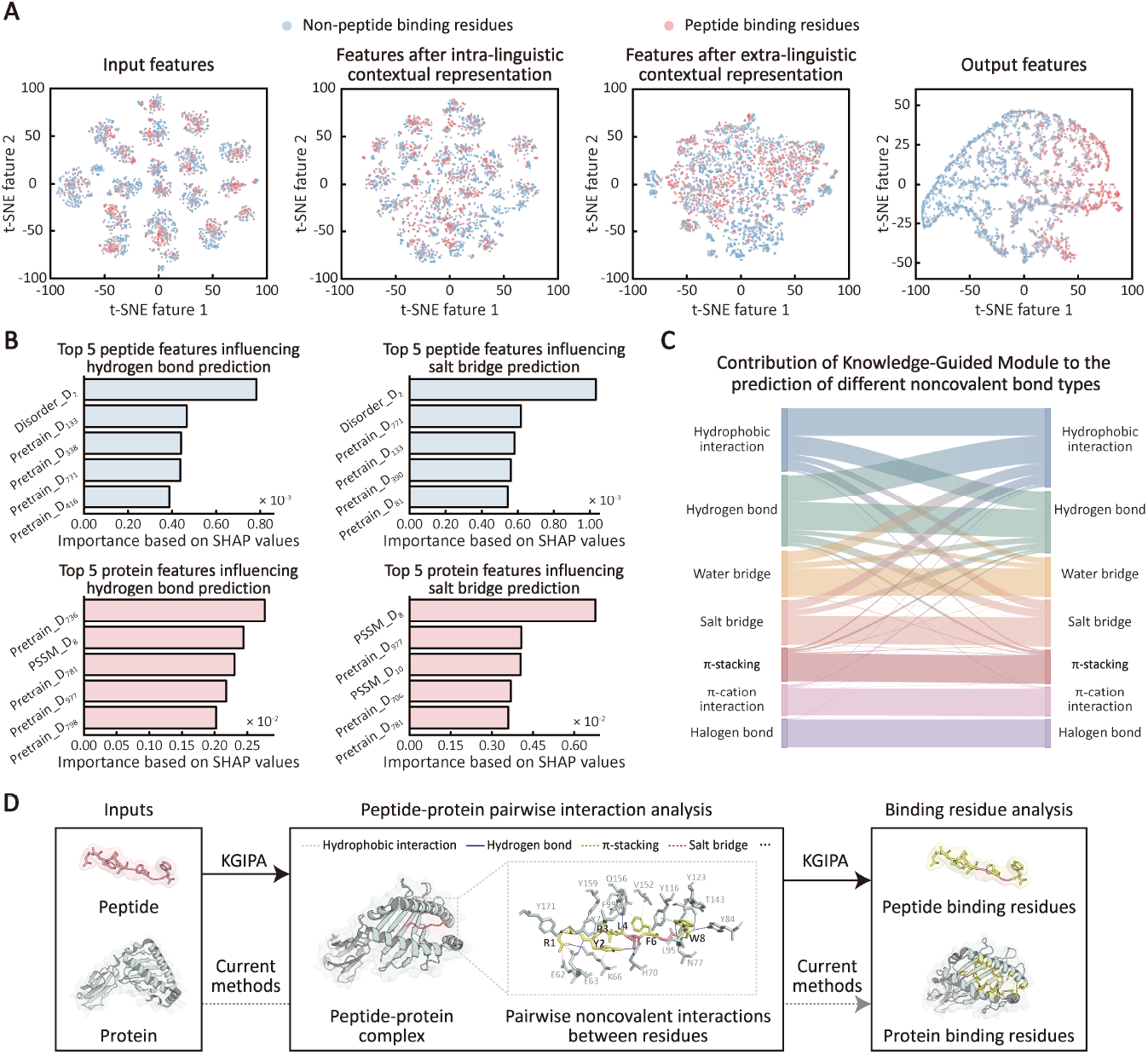
Analysis of KGIPA’s knowledge-guided pragmatic framework and module contributions. (A) t-SNE visualization of feature output in sub-modules based on the independent test dataset Test167. (B) The top 5 peptide and protein features influencing the model’s prediction of hydrogen bond and salt bridge. (C) Contribution of knowledge-guided module to the prediction of different noncovalent bond types. (D) Residue-level interaction maps generated by KGIPA, showing specific peptide-protein residue interactions that provide detailed information beyond conventional binding residue predictions.

To examine the contributions of individual modules, we performed ablation studies (Table 2). The results show that each module—the intra-linguistic contextual representation, the extra-linguistic contextual representation, and the knowledge-guided module—makes a positive contribution in peptide-protein binding site prediction. We further applied SHapley Additive exPlanations (SHAP)^72,73^ to conduct feature-level interpretability analysis, aiming to examine the rationality and effectiveness of the knowledge-guided module (Figure 5B). SHAP is a model-agnostic interpretability framework grounded in cooperative game theory, which explains individual predictions by attributing them to input features through Shapley values^74^. These values quantify each feature’s marginal contribution by averaging over all possible feature coalitions, thereby providing explanations that are theoretically well-founded and uniquely satisfy desirable axioms such as local accuracy, consistency, and missingness^75^. Owing to these properties, SHAP has been widely adopted in bioinformatics to interpret complex machine learning models, substantially facilitating biological insight discovery and advancing the development of the field^76-78^. In this context, SHAP was used to assess the rationality and effectiveness of the knowledge-guided module in KGIPA. The results show that this module adaptively assigns different importance to features depending on the type of non-covalent interaction, while simultaneously capturing common informative patterns across multiple feature categories, including disorder propensity, pre-trained embeddings, and PSSM-based evolutionary features. Overall, these findings demonstrate that the learned feature attributions are not arbitrary but are instead aligned with prior domain knowledge, thereby providing quantitative and theory-backed evidence for the rationality and effectiveness of the proposed knowledge-guided module^17,28^. We also performed Layer-wise Relevance Propagation (LRP)^79^ analysis to examine the contribution of the knowledge-guided module (Figure 5C). By mapping predicted probability scores for each non-covalent bond to the outputs of the extra-linguistic contextual representation, we found that this module consistently contributes positively across all bond types^80^. For example, high mutual contributions between hydrophobic interactions and hydrogen bonds (0.667 for hydrophobic influencing hydrogen bonds, and 0.983 for the reverse) reflect known biological mechanisms, where hydrophobic interactions exclude water molecules and promote aggregation of non-polar groups, thereby facilitating hydrogen bond formation among polar groups. These findings suggest that the knowledge-guided module can autonomously learn associations among different non-covalent bonds without explicit instruction.

**Table 2.** Results of ablation experiments with the KGIPA model.

| Methods | Evaluation Metrics |  |  |  |  |
| --- | --- | --- | --- | --- | --- |
|  | MCC ↑ | ACC ↑ | F1 ↑ | AUROC ↑ | AUPR ↑ |
| **Protein Binding Residue Identification** |  |  |  |  |  |
| KGIPA | **0.552±0.006** | **0.968±0.001** | **0.569±0.005** | **0.941±0.002** | **0.577±0.006** |
| CM1 | 0.518±0.010 | 0.964±0.001 | 0.537±0.010 | 0.933±0.003 | 0.532±0.016 |
| CM2 | 0.526±0.006 | 0.965±0.001 | 0.543±0.006 | 0.935±0.003 | 0.547±0.008 |
| CM3 | 0.539±0.003 | 0.966±0.001 | 0.556±0.004 | 0.939±0.003 | 0.556±0.010 |
| CM4 | 0.538±0.007 | 0.966±0.001 | 0.555±0.007 | 0.941±0.003 | 0.552±0.014 |
| **Peptide Binding Residue Identification** |  |  |  |  |  |
| KGIPA | **0.429±0.005** | **0.738±0.002** | 0.632±0.003 | **0.804±0.003** | **0.694±0.003** |
| CM1 | 0.399±0.004 | 0.702±0.009 | **0.636±0.008** | 0.789±0.002 | 0.667±0.002 |
| CM2 | 0.380±0.011 | 0.686±0.007 | 0.609±0.007 | 0.772±0.006 | 0.611±0.007 |
| CM3 | 0.412±0.009 | 0.712±0.008 | 0.625±0.004 | 0.795±0.006 | 0.677±0.005 |
| CM4 | 0.417±0.009 | 0.717±0.011 | 0.627±0.004 | 0.796±0.004 | 0.676±0.003 |

CM1: Remove intra-linguistic contextual encoder (graph encoder). CM2: Remove extra-linguistic contextual encoder. CM3: Remove knowledge-guided module. CM4: Remove multi-task framework. The results are presented as mean ± standard deviation. The best result for each metric is marked in bold.

At the output level, we examined KGIPA predictions in terms of residue-level interactions. Traditional methods typically report only the set of peptide- or protein-binding residues, without specifying the corresponding interacting residues. By leveraging the extra-linguistic contextual representation, we established a one-to-one correspondence between peptide residues and protein residues, generating residue-level interaction maps that provide detailed insight into molecular interactions. For example, in the 8GVB complex^81^, which involves peptide chain P (an 8-mer peptide from Protein Nef) and protein chain H (MHC class I antigen), conventional methods usually predict binding residues only on one side, such as peptide residues R1, Y2, P3, L4, F6, and W8, without indicating their specific protein partners. In contrast, our maps identify the exact protein residues interacting with each peptide residue (Figure 5D). These detailed interaction maps can guide rational peptide design, support downstream functional studies such as alanine scanning, and help reveal interaction patterns that are difficult to capture with conventional approaches.

### 2.8 Case study on KGIPA predictions

We computed the AUROC and AUPR for KGIPA’s predictions of peptide-protein residue-level pairwise non-covalent interactions on the independent test dataset Test167 (Figure 6A). It is clear that the predictions for these peptide-protein pairs primarily fall into three regions delineated by the average AUROC (value of 0.951) and average AUPR (value of 0.317). Subsequently, we randomly selected a peptide-protein pair from each of the three regions. The AUROC values, AUPR values, and their rankings for the pairwise interaction prediction, peptide- and protein-binding residue prediction are shown in Figure 6B. The true labels and predicted binding residues within the peptide structure and protein sequence are shown in Figures 6C-6E. The first selected peptide-protein pair is from the publicly available TCR TD08 and HLA-A24 complexed with the HIV-1 Nef138-8 peptide (PDB ID: 8GVB)^81^. Among them, the peptide is an 8-mer from Protein Nef (Chain P), and the protein is the MHC class I antigen (Chain H). In this example pair, KGIPA accurately predicted nearly all binding residues of the peptide (with only LEU4 incorrectly predicted as a non-interacting residue), and high prediction scores were noted around the actual interaction residues on the protein sequence. The peptide-protein pair ranked in the middle comes from the human parathyroid hormone receptor-1 dimer (PDB ID: 8HAO)^82^, with the peptide being parathyroid hormone (Chain H) and the protein being the parathyroid hormone/parathyroid hormone-related peptide receptor (Chain I). The predicted binding residues of this peptide show a high overlap with the actual interaction residues (with the recall of 13/16), though a few false positives are also present. A similar phenomenon can be observed on the protein side as well. The third selected peptide-protein pair, with an AUPR ranking around 70% for pairwise non-covalent interactions, is a complex (PDB ID: 7UYK)^83^ composed of RNF31 and FP06655 (a helical peptide). In this pair, the binding residues of the peptide were mostly predicted correctly, with only a few false positives. However, the predicted binding residues in the protein showed a significant deviation from the true labels, likely due to insufficient training data corresponding to this protein. Overall, the visualization results highlight the robust predictive capability of KGIPA and reveal that the prediction difficulty increases as sample imbalance intensifies. Pairwise non-covalent interaction prediction is the most challenging, followed by protein-binding residues, with predicting peptide-binding residues being relatively easier.

**Figure 6.**
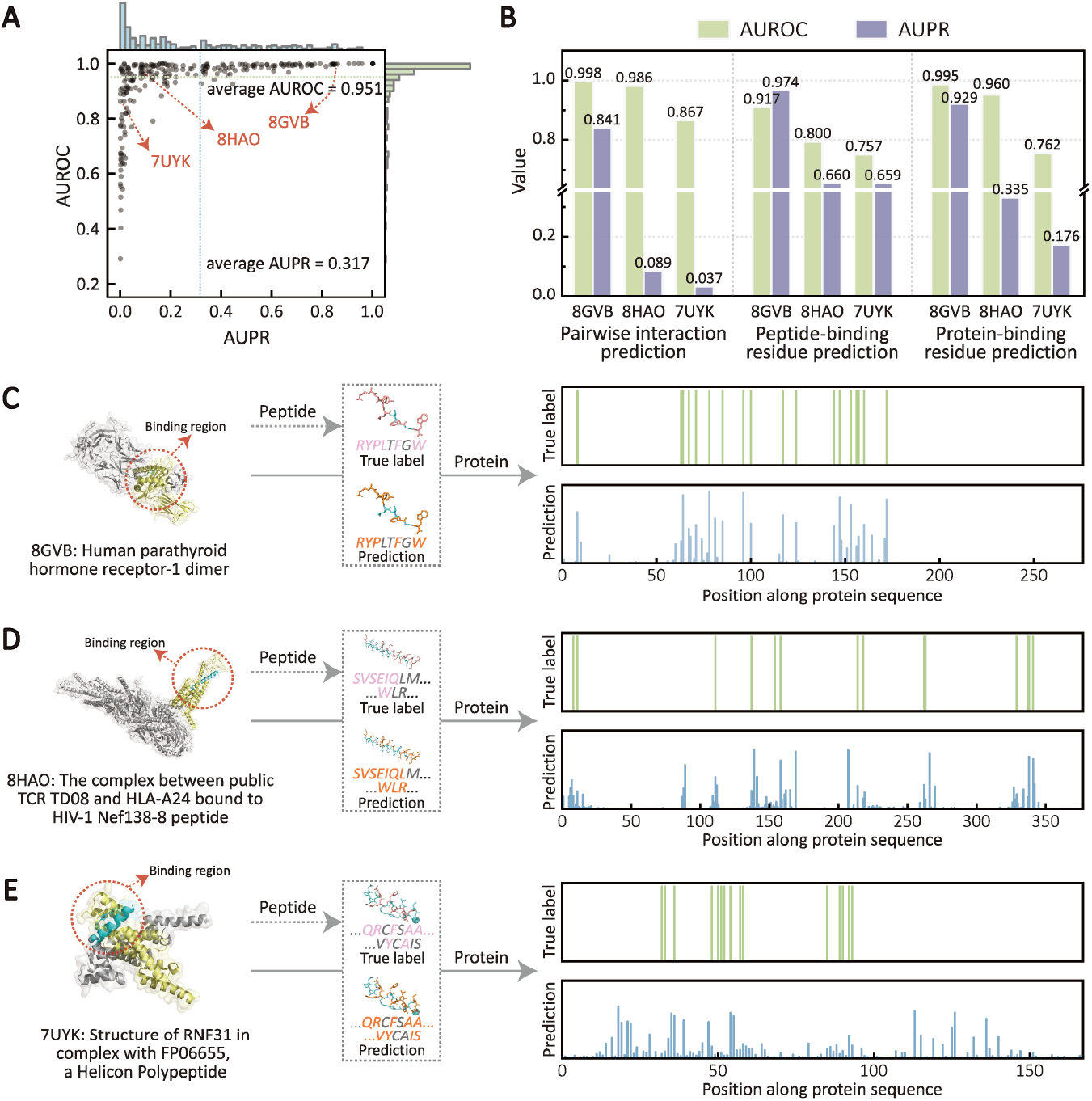
Case studies of three selected examples of peptide- and protein-binding residue predictions. (A) AUROC and AUPR distribution of KGIPA pairwise prediction results based on the independent test dataset Test167. (B) AUROC and AUPR values for pairwise interaction and binding residue prediction across three selected pairs. (C) Analysis of KGIPA prediction results based on complex 8GVB. (D) Analysis of KGIPA prediction results based on complex 8HAO. (E) Analysis of KGIPA prediction results based on complex 7UYK.

### 2.9 Computational efficiency analysis of KGIPA

To better demonstrate the computational performance of our method, we conducted an efficiency analysis comparing KGIPA with representative baseline models. Detailed results and analysis are provided in Note S2 and Table S6. The analysis shows that the primary computational cost of KGIPA arises from the molecular representation stage, particularly from the prediction of 3D structural features using trRosetta and the prediction of two-dimensional structural features using SCRATCH, as well as the modeling of pairwise residue-level interaction graphs in the extra-linguistic context module. While this increases training time compared with baseline methods, the inference time for new data is comparable, allowing KGIPA to achieve high predictive performance without incurring additional overhead during practical use.

Furthermore, considering the scalability and flexibility, we developed a parallelized variant of KGIPA called KGIPA-Fast by removing the 3D structural features derived from trRosetta and the secondary structural features obtained from SCRATCH. This variant substantially reduces preprocessing and training time, with the average AUPR decreasing slightly from 0.633 to 0.607, which remains competitive with existing methods. Researchers can flexibly choose KGIPA or KGIPA-Fast according to their application scenarios and specific task requirements. In addition, for high-throughput computations, users can deploy the standalone version of KGIPA locally to perform parallel processing or model retraining. These results indicate that KGIPA not only exhibits strong predictive capability but also provides a scalable and flexible deployment solution for diverse research contexts.

### 2.10 KGIPA introduces a new computational framework for biological sequence analysis

At the core of KGIPA lies the biological sequence pragmatic analysis, which integrates intrinsic and extrinsic factors that affect sequence functions by representing the intra- and extra-linguistic context. Compared to existing methods that simply concatenate multiple features, KGIPA constructs a structured pipeline from “intra-linguistic contextual representation” to “extra-linguistic contextual representation” and ultimately to “pragmatic analysis”, enabling a deeper understanding and profiling of actual biological processes. In bioinformatics, numerous tasks are related to or share similarities with peptide-protein interaction prediction, such as peptide-protein binding affinity prediction, protein-protein interaction (PPI) prediction, drug-target interaction (DTI) prediction, and drug-disease association prediction^84^. Consequently, we further explored whether the proposed biological sequence pragmatic analysis pipeline could demonstrate generalizability and offer advantages in similar tasks.

Specifically, we adapted and retrained the KGIPA model using the peptide-protein binding affinity prediction task as an example. Predicting the binding affinity between peptides and proteins is crucial for understanding physiological and pathological cellular processes, which in turn facilitates the identification of potential therapeutic candidates and the development of personalized treatment strategies. Given the regression property of binding affinity prediction, the core structure of KGIPA—representing both intra- and extra-linguistic context to realize pragmatic analysis—was preserved during adaptation. Only the decoder, composed of fully connected layers, was modified, and the multi-task learning framework irrelevant to affinity prediction was removed. Additionally, it is important to note that the peptide-protein binding affinity prediction data was sourced from the PDBbind v2020 database and preprocessed using our constructed feature library. Experimental results indicate that KGIPA outperformed other advanced prediction methods, excelling in multiple evaluation metrics such as root mean square error (RMSE), mean absolute error (MAE), coefficient of determination (R^2^), and Pearson correlation coefficient. Furthermore, results from cross-validation experiments demonstrated that KGIPA exhibits better stability (Figure 7A).

**Figure 7.**
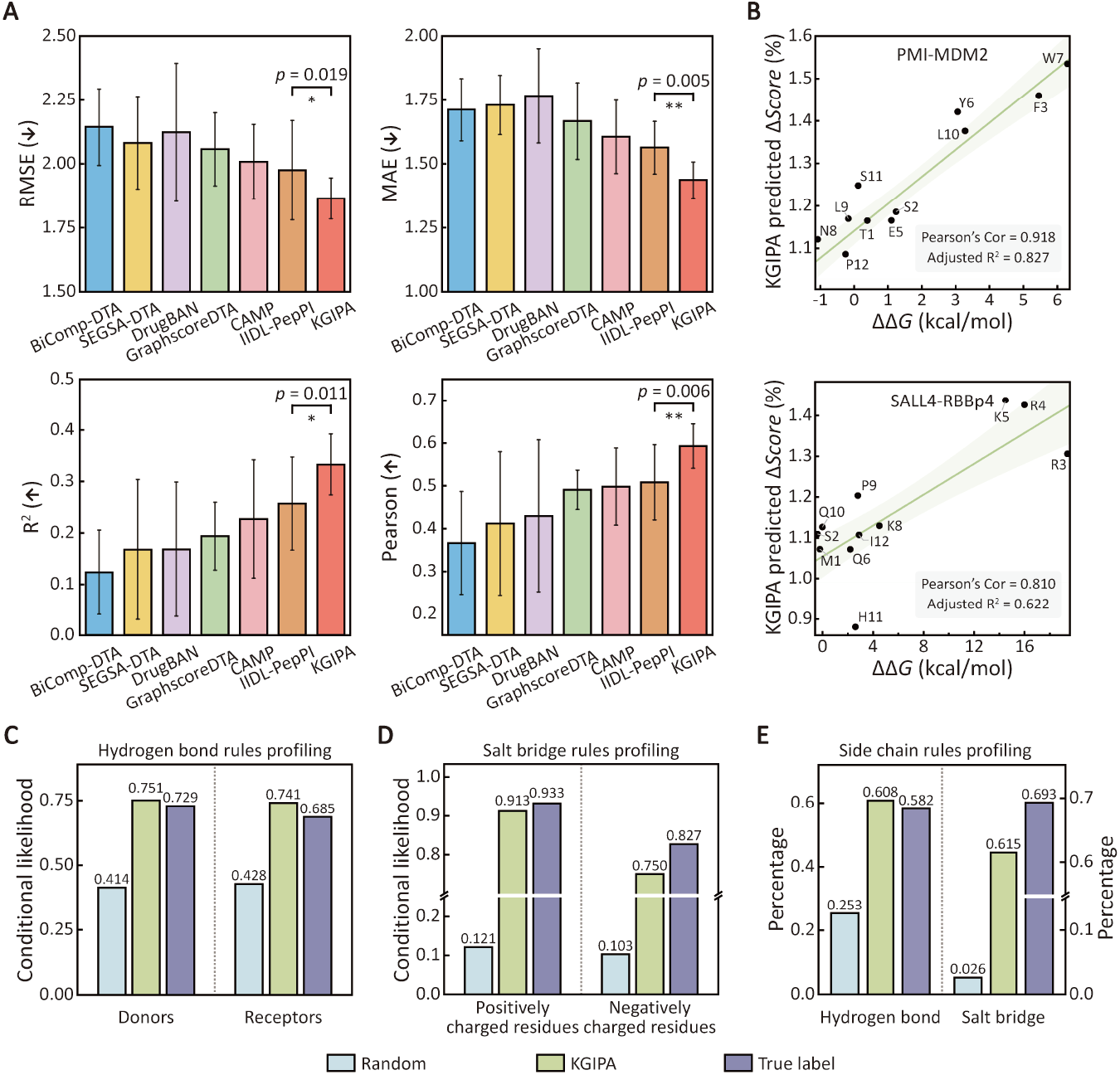
Experimental results of KGIPA in peptide-protein binding affinity prediction and two related extended application tasks. (A) Comparison of KGIPA’s performance with other advanced binding affinity prediction methods. (B) For the peptide alanine mutants of PMI and SALL4, comparisons between the interaction capability changes predicted by KGIPA and the experimentally measured changes in Gibbs free binding energy. (C) Conditional likelihood scores for hydrogen bond donors and acceptors. (D) Conditional likelihood scores for positively and negatively charged residues in salt bridges. (E) Analysis of common bonding rules for side-chain hydrogen bonds and salt bridges.

In summary, the biological sequence pragmatic analysis pipeline proposed by KGIPA has demonstrated good generalizability and high accuracy in peptide-protein interaction and binding affinity prediction tasks. Given the complexity of data collection and curation for different tasks, although we have not conducted further tests and evaluations on a broader range of tasks, KGIPA at least offers a new computational framework for AI-driven life science research, serving as a viable approach to address critical challenges in biological sequence analysis and association prediction.

### 2.11 Extended application of KGIPA in two related tasks

#### 2.11.1 KGIPA facilitates key residue identification for peptide drug design

Unlike binding residues in peptide-protein non-covalent interactions, which reflect their binding capabilities with specific biomolecules, key residues are crucial for the function of the peptide or protein. They may impact structural stability and overall function^85^. Therefore, the relationship between binding and key residues is not always straightforward and requires further experimental validation and functional analysis^86^. To assess KGIPA’s potential in identifying key residues and guiding peptide drug design, we employed a comparative approach involving dry- and wet-lab experiments to evaluate the consistency between KGIPA’s predictions and biological assays. Specifically, we employed the alanine scanning strategy^87^ to systematically replace the amino acid residues in the peptide and calculate the sum of non-covalent interaction probabilities for each mutation at the residue level. This was done to evaluate changes in the interaction capability of each peptide mutant with a given protein (Δ*Score* = (1 + (*Score*_*WT*_ - *Score*_*Ala*_) / *Score*_*WT*_)), where *Score*_*WT*_ denotes the interaction probability of the wild-type peptide, and *Score*_*Ala*_ denotes that of the alanine mutant peptide. Then, we can identify the key residues driving peptide function based on the Δ*Score* and compare these with changes in Gibbs free binding energy (ΔΔ*G*) from wet-lab experiments to explore the feasibility of using KGIPA to guide peptide drug design. Based on the principles above, we explored KGIPA’s predictions of key residues using two peptide-protein pair examples: PMI (a potent, synthetically produced 12-mer peptide inhibitor) with the Oncoprotein Mouse Double Minute 2 Homolog (MDM2)^88^, and Sal-like 4 (SALL4) with the Retinoblastoma Binding Protein 4 (RBBp4)^89^. The experimental results are shown in Figure 7B. Notably, in the PMI-MDM2 interaction, KGIPA predicted that alanine substitutions at residues Phe3, Tyr6, Trp7, and Leu10 on the peptide side would significantly reduce the interaction capability of the mutant peptides with MDM2. This is consistent with the key residues identified through wet-lab experiments (Pearson correlation coefficient = 0.918, adjusted R^2^ = 0.827, for details, see Note S3 and Figure S3)^90^.

Similarly, for the SALL4-RBBp4 interaction, we observed results consistent with wet-lab experiments, identifying Arg3, Arg4, and Lys5 as the key residues mediating peptide function (Pearson correlation coefficient = 0.810, adjusted R^2^ = 0.622)^91^. These findings demonstrate that using KGIPA to identify key residues in peptide-protein interactions is feasible, yielding results highly consistent with wet-lab experiments while shortening analysis time, which helps accelerate peptide drug development.

To further support KGIPA’s context sensitivity, we conducted additional analyses, including systematic alanine scanning of the same peptide across different protein partners^92-94^, combined computational-experimental validation of predicted key residues^95-98^, and evaluation of phosphorylation effects^99^. Detailed descriptions of these experiments are provided in Note S3, Figures S4-S5 and Table S8. Overall, these analyses demonstrate that KGIPA reliably captures context-dependent variations in peptide-protein interactions and accurately identifies key residues under different receptor environments, supporting its effectiveness in modeling biologically relevant context dependence.

#### 2.11.2 KGIPA for revealing the non-covalent rules of peptide-protein interactions

To explore the underlying biological mechanisms, we investigated whether KGIPA could capture the non-covalent rules of peptide-protein interactions. Specifically, within the collected training dataset, we first quantified the number and proportion of different types of non-covalent bonds, as shown in Table S9. Due to the presence of hydrophobic carbon atoms in the 20 standard amino acids, hydrophobic interactions have the highest proportion in the data and exhibit highly complex relationships, making them challenging to analyze^25^. Similarly, water bridges often serve as auxiliary interactions to other non-covalent bonds, and their formation is highly dependent on the surrounding water environment, further complicating direct analysis^100^. Therefore, we focused our analysis on the chemical rules for forming hydrogen bonds and salt bridges, which are more tractable and crucial to peptide-protein interactions.

We aimed to describe the bonding rules of protein residues with specific properties based on the properties of their interacting peptide residues, following the approach of Li *et al*.^25^. The conditional likelihood score is defined as: *p* (protein residue property = *x* | peptide residue property = *y*) = (number of protein residues ∈ *S*(*x*) that form specific non-covalent bonds with peptide residues of property *y*) / (total number of protein residues that form specific non-covalent bonds with peptide residues of property *y*). Here *S*(*x*) denotes the set of residues with side chains that contain at least one element meeting property *x*. For example, *S*(“hydrogen bond donor”)={H, K, N, Q, R, S, T, W, Y, C}, which includes residues that have at least one hydrogen bond donor in their side chains. Similarly, *S*(“hydrogen bond acceptor”)={M, W, D, E, H, N, Q, S, T, Y}, *S*(“salt bridge positive residue”)={K, R, H}, and *S*(“salt bridge negative residue”)={D, E}. Using these definitions, we calculated the conditional likelihood scores for hydrogen bonds and salt bridges in three scenarios—random selections (used as the control), KGIPA predictions, and true labels.

First, we analyzed hydrogen bonds among non-covalent interactions. Hydrogen bonds typically form between a hydrogen donor and an acceptor group. When peptide residues act as hydrogen acceptors, the conditional likelihood score for hydrogen bond donor residues in proteins as interaction partners (0.729, true labels) is significantly higher than for control residues (0.414, random selections). KGIPA’s predicted score for hydrogen bond donor residues in proteins is also high and closely matches the true labels (0.751, see Figure 7C). Similarly, when peptide residues serve as hydrogen bond donors, the likelihood score for protein acceptor residues predicted by KGIPA is notably higher than the control and aligns closely with the true labels. Additionally, salt bridges are stable interactions formed between oppositely charged amino acid residues through electrostatic forces. When peptide residues carry a negative charge, the conditional likelihood score for positively charged residues in proteins (0.933, true labels) is much higher than for random controls (0.121). KGIPA’s predicted scores for salt bridge sites also exceed random controls and are closely aligned with the true label results (Figure 7D). A similar phenomenon is observed when peptide residues act as positive charge residues of salt bridges, further demonstrating KGIPA’s capability to learn the non-covalent rules of peptide-protein interactions.

Furthermore, we analyzed the proportion of side-chain hydrogen bonds adhering to common bonding rules. The KGIPA prediction results closely resemble those based on true labels (Figure 7E). Similarly, the proportion of side-chain salt bridges following common bonding rules was calculated, and these results align with our previous findings. In summary, the results indicate that KGIPA effectively captures the non-covalent bonding rules of peptide-protein interactions, which aids researchers in uncovering unknown or complex non-covalent mechanisms.

## 3 DISCUSSION

In this study, we introduce KGIPA, an end-to-end deep learning approach designed to predict pairwise non-covalent interactions between peptides and proteins while also identifying the types of non-covalent bonds involved. During the modeling process, we fully integrate information on peptide-protein pairs through intra- and extra-linguistic contextual representations and enhance feature interactions between different bonding types through the knowledge-guided module, thereby realizing biological sequence pragmatic analysis. Experimental results demonstrate that KGIPA surpasses other state-of-the-art methods in non-covalent interaction prediction and bond type identification. In addition, by leveraging the features learned through intra- and extra-linguistic contextual representations and the knowledge-guided module, KGIPA can provide biologically relevant insights into the predicted interactions. The biological sequence pragmatic analysis pipeline proposed by KGIPA demonstrates strong generalizability and can be easily extended to other related fields or similar tasks, providing a new computational approach for AI-driven life science research.

This work focused on integrating multi-modal features of peptides and proteins, including predicted 3D structures, to more accurately represent intra-linguistic contexts. By combining intra- and extra-linguistic contextual representations, KGIPA achieves biological sequence pragmatic analysis, enabling residue-level pairwise non-covalent interaction predictions. Moving forward, KGIPA can be extended in several ways. First, current peptide-protein interaction prediction methods are limited to the dimer level, meaning they only support the analysis of individual peptide-protein pairs. However, in actual complexes, multiple peptide-

protein pairs coexist and influence each other^101^. Therefore, extending KGIPA’s pragmatic analysis approach to investigate interactions in complex multimers assemblies is an exciting direction for future exploration. Second, although KGIPA provides residue-level pairwise interaction predictions, pharmacologists developing new drugs often prefer to obtain the complete 3D structure of a complex. Thus, refining residue-level predictions to atomic-level 3D spatial structures could better facilitate peptide drug development and protein function research. Finally, KGIPA’s framework and residue-level interaction modeling provide a strong foundation for rational peptide design^102,103^. By identifying key binding residues and their contextual dependencies, KGIPA can guide the optimization of peptide sequences for enhanced affinity, specificity, and stability, thereby facilitating therapeutic development.

## 4 METHODS

### 4.1 Data collection and label extraction

In this study, we meticulously curated non-covalent interaction data from all peptide-protein complexes in the RCSB Protein Data Bank (PDB) database before July 2024. Specifically, the rule-based tool PDB-BRE was employed to identify all complexes involving peptide-protein interactions within the database. To reduce redundancy while retaining diverse coverage of the data, we applied CD-HIT to remove peptide-protein pairs in the training dataset that shared more than 80% sequence similarity with those in the test datasets (LEADS-PEP, Test251, Test167, and Test1440). The sequence similarity threshold is recommended and widely adopted in related studies^7,50^. To further assess model generalization, we additionally constructed independent test subsets using more stringent 60% and 40% sequence similarity thresholds. In addition, to ensure data quality, we excluded peptide-protein pairs in which more than 20% of the residues were unknown or non-standard amino acids.

Subsequently, we employed PLIP to annotate residue-level non-covalent interactions, covering seven interaction types, including hydrophobic interactions, hydrogen bonds, water bridges, salt bridges, π stacks, π-cation interactions, and halogen bonds^32^. Because our study focuses on identifying interaction residues in peptide-protein pairs, positive and negative samples were defined at the residue level: interaction residues were regarded as positive samples, whereas non-interaction residues were considered negative samples.

On the peptide side, about 32.6% of residues were involved in interactions, largely due to the short length of peptides, which often results in a substantial fraction of residues directly participating in binding. In contrast, only about 3.7% of protein residues were labeled as positives, reflecting the much greater length of protein sequences, in which only a small subset of surface or functional-site residues typically mediate binding. Detailed statistical information for the datasets is provided in Tables S1 and S9. The peptide-protein interaction data collected in this study are accessible via the online prediction webserver.

### 4.2 KGIPA model architecture

#### 4.2.1 Intra-linguistic contextual representation

For the sequence features of peptides and proteins, we utilized a variety of tools, including integer encoding, physicochemical property encoding, PSI-BLAST^38^, IUPred2A^39^, SCRATCH^40^, and ProtT5^17^, to represent the amino acid position, physicochemical properties, evolutionary information, disorder tendencies, secondary structure, and pre-trained embeddings. To ensure fairness and compatibility across multiple sources, we unified the dimensions of these features using word embeddings or linear transformations. As for the structural features, directly extracting them from complexes may lead to information leakage, which is not conducive to analyzing novel peptides and proteins. Therefore, based on the original sequences, we employed trRosetta^41^ to predict 3D structures and extract the spatial relationships between residues. The contact map predicted by trRosetta includes distance and angle information for all residue pairs. In this study, we focus on the distance information between pairs of Cβ atoms in the contact map to illustrate the spatial relationships among residues. The distance information denoted as **A** ∈ℝ^*l*×*l*^, is a binary matrix where *l* represents the total number of amino acids, and each element *a*_*uv*_ is defined as^104^:

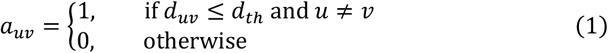

where *d*_*uv*_ represents the distance between the *u*-th and *v*-th atoms, and *d*_*th*_ is the threshold distance. Following the study by Wang *et al*.^42^, we set *d*_*th*_ to 20 in this work.

To streamline feature input and model training, we introduced a preprocessing step to standardize the lengths of both peptide and protein sequences. Specifically, peptides shorter than 50 residues were padded with the letter “X” to ensure uniform length. Similarly, proteins shorter than 800 residues were padded with “X” to reach a length of 800^7^. For proteins longer than 800 residues, we applied the truncation method proposed by Zhang *et al*.^105^, maintaining a maximum length of 800.

For each peptide or protein, the sequence in Fasta format is converted into a graph representation *G*={*V, E*} through the feature mentioned above extraction process. In this graph, the nodes *V* represent the sequence residues, and the edges *E* correspond to their contact relationships. To ensure compatibility, we apply word embeddings or linear transformations to obtain a dense node feature matrix **V**∈ ℝ^*L*×*D*^, where *L* is the predefined maximum number of nodes (equal to the sequence length), and *D* represents the feature dimension for each node. Specifically, the node feature matrices for all proteins (denoted *M* proteins) are organized as a set **R** = {**R**_**1**,_ …, **R**_*i*_, …, **R**_*M*_}, where 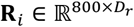is the node feature matrix for the *i*-th protein, and *D*_*r*_ indicates the feature dimension for proteins. Similarly, the node feature matrices for all peptides are expressed as **E** ={**E**_1,_ …, **E**_*i*_, …, **E**_*N*_}, where *N* is the total number of peptides, and the node feature matrix for the *j*-th peptide is given by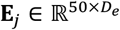, with *D*_*e*_ denoting the feature dimension for peptides.

We employ two independent two-layer GCN modules to effectively learn the intra-linguistic contextual representations of peptides and proteins, respectively. The GCN updates node feature vectors by aggregating neighboring residues linked by peptide bonds, extending convolution operations to irregular domains to capture 3D structural information. To address issues of vanishing or exploding gradients during deep network training, residual connections are applied between layers of the GCN modules. For the output, we retain the node-level representations for both peptides and proteins. Therefore, taking the *i*-th protein as an example, the intra-linguistic context encoder can be expressed as:

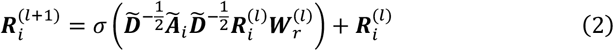

where Ã_*i*_ is the adjacency matrix of protein *i* with self-loops added, and 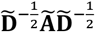denotes the normalization operation on the adjacency matrix. 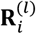is the input node feature matrix of the *i*-th protein at layer *l* (note that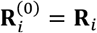),is the output node feature matrix for the *i*-th protein at layer *l*+1. 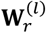is the learnable weight matrix for the *l*-th layer GCN used for protein intra-linguistic contextual representation, and *σ*(·) denotes a non-linear activation function, with ReLU(·) used in our model.

Similarly, the intra-linguistic context encoder for the *j*-th peptide representation can be denoted as follows:

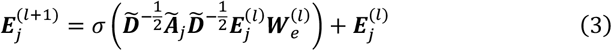

where 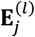 is the input node feature matrix of peptide *j* at layer *l* (note that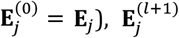is the output node feature matrix for the *j*-th peptide at layer *l*+1, and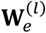is the learnable weight matrix for the *l*-th layer GCN used for peptide intra-linguistic contextual representation.

Next, we map the intra-linguistic contextual representations of peptides and proteins to generate representations specific to various non-covalent bonds. Specifically, for peptides (or proteins), we designed fully connected layers with a consistent output dimension for each of the seven non-covalent bond types, incorporating dropout regularization to mitigate overfitting. Taking hydrophobic interactions as an example, the intra-linguistic contextual representation of the *i*-th protein, after being mapped through the fully connected layer, can be expressed as:

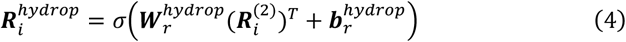

where 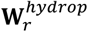and 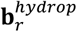correspond to the weight matrix and bias vector used to represent hydrophobic interactions in proteins, 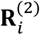denotes the representation of the *i*-th protein generated by the GCN encoder in the second layer, while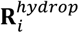 represents the *i*-th protein mapped specifically to hydrophobic interactions. Similarly, representations for other non-covalent bonds can be obtained for proteins and peptides.

#### 4.2.2 Extra-linguistic contextual representation

We utilize bilinear interaction-based encoders for extra-linguistic contextual representation to effectively capture the pairwise non-covalent interactions that occur between peptides and proteins. For example, when analyzing hydrophobic interactions, the representations of specific non-covalent bonds obtained by mapping for protein *i* and peptide *j* can be expressed as 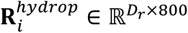and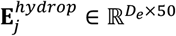. To facilitate the construction of bilinear interaction map, we first transform these representations to ensure compatible feature dimensions, which can be expressed as:

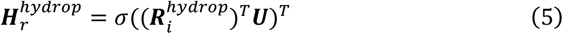

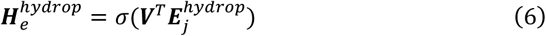

Where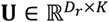and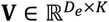are learnable weight matrices for transforming the dimensions of proteins and peptides, respectively. We then use these updated representations to construct the bilinear interaction map, resulting in a matrix 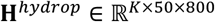that reflects the hydrophobic interactions:

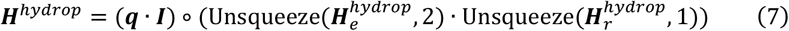

where **q** ∈ ℝ^*K*^ is a learnable weight vector, **I** ∈ ℝ^50×800^ is a fixed all-ones matrix and ° denotes Hadamard (element-wise) product. Therefore, the submatrix **h**_*k*_ in **H**^*hydrop*^ represents the *k*-th feature reflecting the strength of the peptide-protein hydrophobic interaction. To better illustrate the bilinear interaction, the submatrix **h**_*k*_ in equation (7) can also be expressed as:

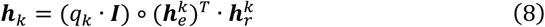

Where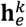is the *k*-th row of 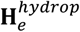and 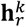is the *k*-th row of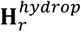,denoting the *k*-th feature representation of the peptide and protein, respectively. *q*_*k*_ is the *k*-th value of **q**.

Additionally, since padding was applied to the peptide and protein sequences during feature extraction, we further applied a masking operation to the non-covalent bond representations. This eliminates the noise in the representation matrices and helps accelerate model training. The masking process is as follows:

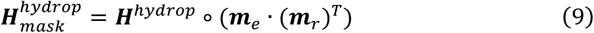

where **m**_*e*_ is the mask vector for the sequence length of peptide *j* (1 for the presence of amino acids, 0 otherwise) and **m**_*r*_ is the mask vector for protein *i*. Therefore, using extra-linguistic contextual representations similar to those used for hydrophobic interactions, we can obtain representations for the other six types of non-covalent bonds: hydrogen bonds, water bridges, salt bridges, π stacks, π-cation interactions, and halogen bonds.

Non-covalent interactions between peptides and proteins are mediated by various non-covalent bonds, which exhibit interdependencies and correlations. Different non-covalent bonds often work together to either facilitate or hinder specific binding modes, thus affecting the stability and functionality of the complex^106^. For example, an excess of hydrophobic interactions can alter the polarity environment of the binding site, potentially reducing the likelihood of forming hydrogen bonds or salt bridges. Therefore, integrating representations 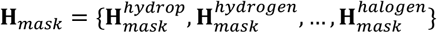obtained from extra-linguistic contextual representations of different non-covalent bonds will help the model learn potential correlations among these bonds. Specifically, we first performed a statistical analysis on the collected data, extracting prior knowledge about the correlations between different non-covalent bonds by calculating correlation coefficients and representing this knowledge with an adjacency matrix **T**∈ ℝ^7×7^. Then, we integrated the representations of different non-covalent bonds to learn the underlying associations:

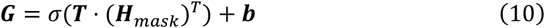

where **b**∈ ℝ^7^ is the learnable bias vector for the knowledge-guided module.

Next, the interaction strength representations for different non-covalent bonds are fed into fully connected layers with a sigmoid activation function to obtain pairwise prediction probabilities for each bond type. The prediction process can be expressed as:

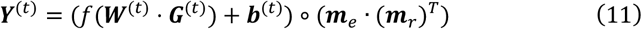

where **W**^(*t*)^ and b^(*t*)^ represent the learnable weights and bias in the fully connected layer for predicting the type *n* non-covalent bond, **G**^(*t*)^ denotes the pairwise representation for the *t*-th non-covalent bond, and *f* (·) is the sigmoid activation function. After passing through fully connected layers, the prediction results are multiplied by the mask matrix (**m**_e_ · (**m**_*r*_)^T^) to limit the effective range of pairwise predictions. Thus, we concatenate the prediction results along the non-covalent bond type dimension to form a new three-dimensional tensor **Y**^**TNCI**^={**Y**^(0)^…,**Y**^(*t*)^,…,**Y**^(6)^}∈^7 ×50×800^.

Furthermore, global max pooling can be applied to the above prediction results for the seven non-covalent bonds, thereby obtaining peptide-protein non-covalent interaction predictions **Y**^NCI^ without distinguishing bond types.

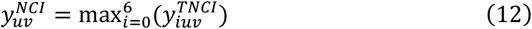

where 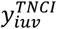represents the predicted probability of the *i*-th type of non-covalent bond existing between residue pairs *u*-*v*, and 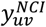denotes the non-covalent interaction probability between residue pairs *u*-*v*.

#### 4.2.3 Biological sequence pragmatic analysis

KGIPA adopts the uncertainty-weighted loss function^107^ based on maximum Gaussian likelihood estimation to achieve joint prediction of peptide-protein non-covalent interactions and non-covalent bond types. This function adjusts the loss for each task according to the associated uncertainty, allowing the model to balance performance across different tasks more effectively. It can be expressed as:

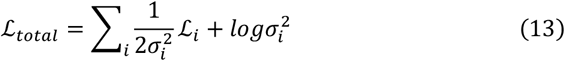

where *i* denotes the task index, 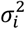represents the uncertainty estimation for the *i*-th task, and 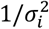is the uncertainty weight, which determines the contribution of each task’s loss to the overall loss. The term 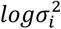serves as a regularization term to penalize the increase in uncertainty estimation, preventing the model from minimizing the overall loss by infinitely increasing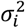. Each subtask can be treated as a binary classification task, and thus, based on the binary cross-entropy loss function, ℒ_*i*_ can be expressed as:

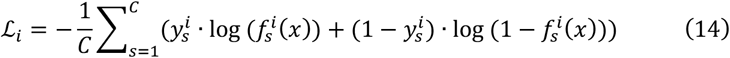

where *C* is the number of peptide-protein residue pair samples, 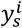represents the actual label of the *s*-th residue pair in the *i*-th subtask, and 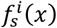represents the predicted probability of the *s*-th residue pair in the *i*-th subtask.

## RESOURCE AVAILABILITY

### Materials availability

This study did not generate new materials.

### Data and code availability

- The experimental data supporting this study are available in the Zenodo repository (https://doi.org/10.5281/zenodo.17383805) and via the KGIPA online prediction webserver (http://bliulab.net/KGIPA/document). All data used in this work are from public resources. The structural information of the peptide-protein complexes used in this work can be downloaded from the RCSB PDB database (https://www.rcsb.org/), the peptide-protein interactions in the complexes can be accessed via PDB-BRE (http://bliulab.net/PDB-BRE), and the information on the paired non-covalent binding residues can be obtained from PLIP (https://plip-tool.biotec.tu-dresden.de/plip-web/plip/). The independent test datasets are available from Test167 and Test1440 (https://doi.org/10.5281/zenodo.17383805), LEADS-PEP (https://pubs.acs.org/doi/full/10.1021/acs.jcim.5b00234), Test251 (https://bitbucket.org/isaakh94/interpep_pipeline/src/master/databases/). All the data used in this work can be obtained from the corresponding authors.
- The KGIPA and KGIPA-Fast online prediction webserver is detailed at http://bliulab.net/KGIPA, while the standalone package developed in this study can be freely accessed on GitHub at https://github.com/ShutaoChen97/KGIPA.

## ACKNOWLEDGMENTS

This work was supported by the National Natural Science Foundation of China (No. 62325202, U22A2039).

## AUTHOR CONTRIBUTIONS

S.C., K.Y., and B.L. conceived the concept. S.C. designed methodology and performed experiments. K.Y. and J.S. assisted in the experiments. S.C. wrote the manuscript. S.C., K.Y., X.Z. and B.L. contributed to the revision of the manuscript. B.L. supervised this work.

