## Supplementary Information for "Pragmatic analysis with knowledge-guided for unraveling peptide-protein pairwise non-covalent mechanisms"

### Supplementary Figures

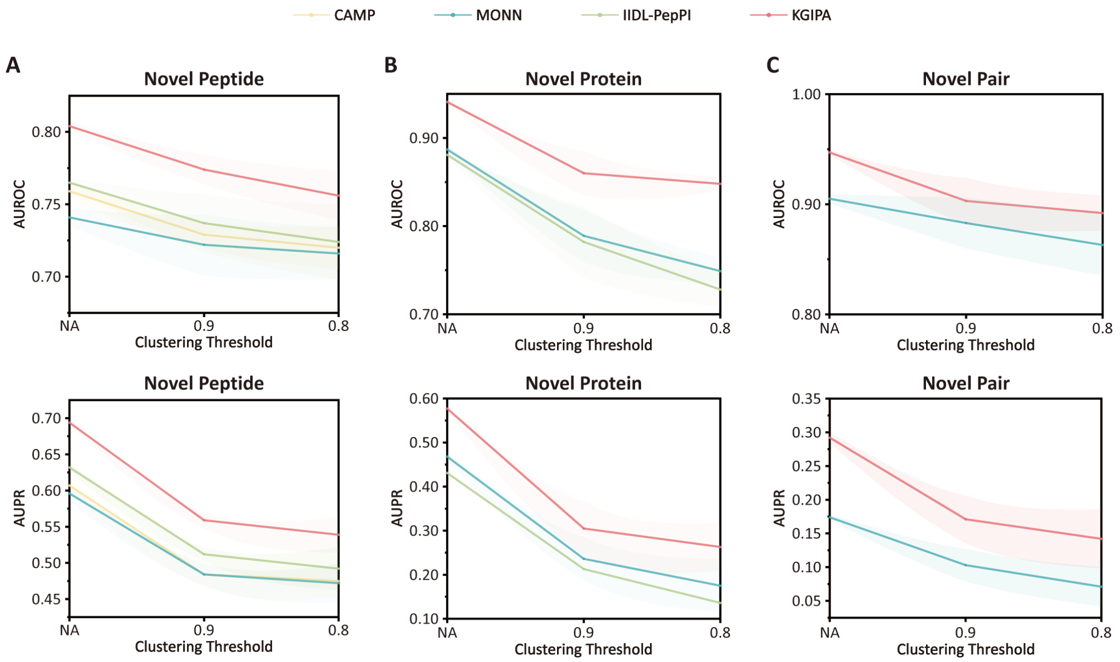


**Supplemental Figure 1: Generalization performance evaluation of KGIPA and existing methods under three scenarios.** (**A**) novel peptides, (**B**) novel proteins, and (**C**) novel peptide-protein pairs.

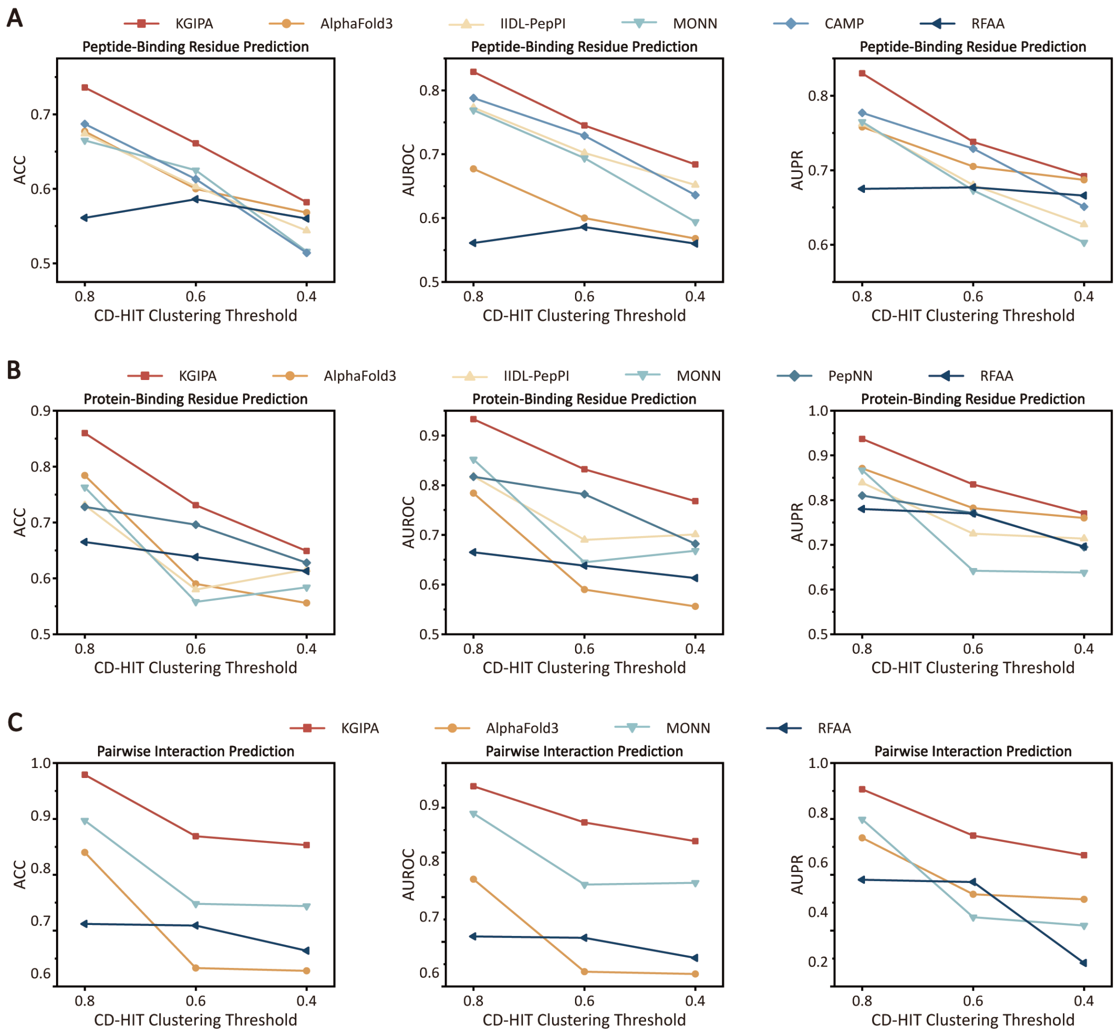


**Supplemental Figure 2: Performance of KGIPA and state-of-the-art methods under different CD-HIT sequence similarity thresholds.** (**A**) Peptide-binding residue prediction under different thresholds. (**B**) Protein-binding residue prediction under different thresholds. (**C**) Pairwise interaction prediction under different thresholds.

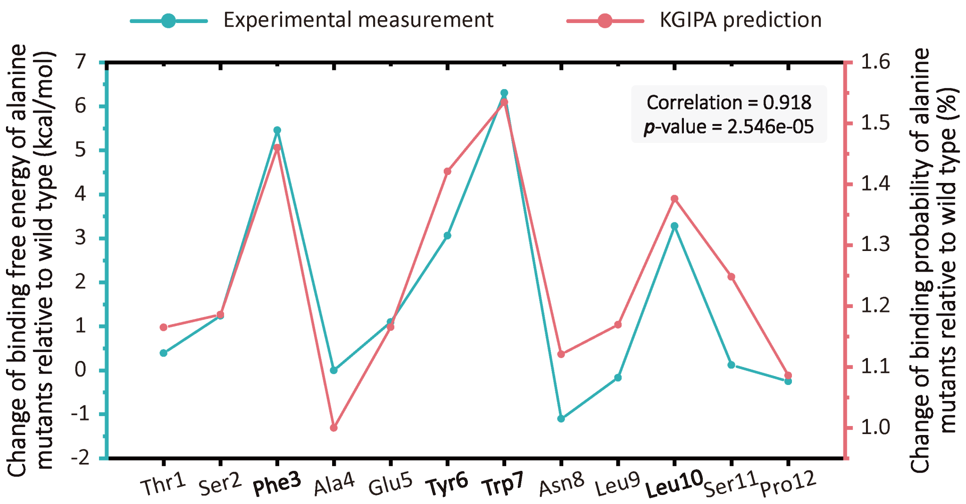


**Supplemental Figure 3: Relative change in the binding probability of MDM2 with PMI peptide alanine mutants predicted by KGIPA versus experimentally measured changes in binding free energy.**

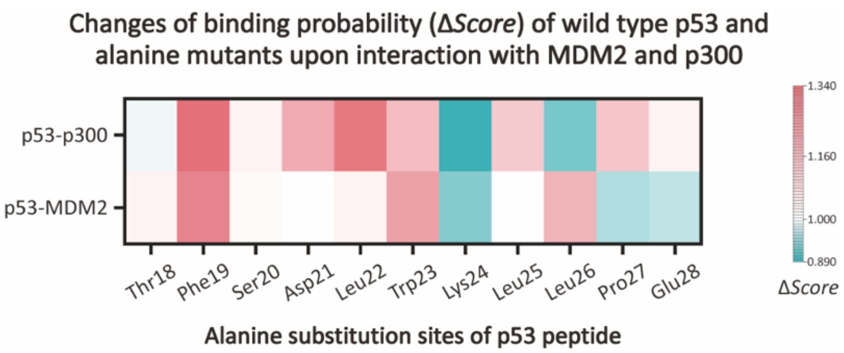


**Supplemental Figure 4: Heatmap of KGIPA-predicted binding probability changes (Δ*Score*) for wild-type and alanine-substituted p53 peptides interacting with MDM2 and p300.**

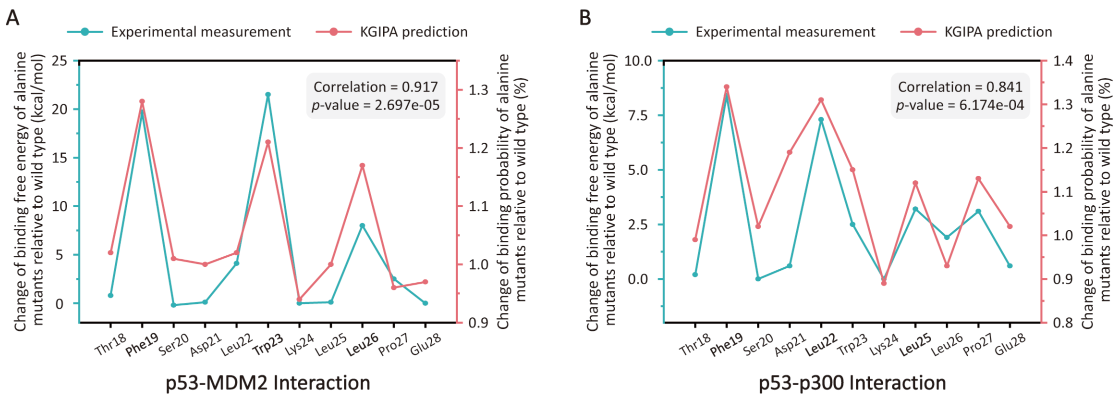


**Supplemental Figure 5: Comparison of KGIPA-predicted key residues with experimentally determined ΔΔ*G*.** (**A**) p53-MDM2 interaction. (**B**) p53-p300 interaction.

#### Supplementary Tables

**Supplemental Table 1. Detailed of the peptide-protein interaction datasets used in this study.**

| Dataset | Training dataset | Test251 | LEADS-PEP | Test167 | Test1440 |
|---|---|---|---|---|---|
| **Pairs** | 8,622 | 250 | 52 | 246 | 1,440 |
| **Peptides** | 6,313 | 239 | 50 | 166 | 886 |
| **Proteins** | 5,688 | 250 | 52 | 175 | 937 |
| **Peptide binding residues, n (%)** | 52,530 (32.66%) | 1,795 (48.55%) | 306 (69.07%) | 1,438 (29.32%) | 8,503 (26.72%) |
| **Protein binding residues, n (%)** | 85,594 (3.72%) | 2,841 (4.58%) | 556 (5.72%) | 2,394 (3.09%) | 13,749 (3.13%) |
| **Pairwise interacting residues, n (%)** | 105,030 (0.24%) | 3,577 (0.39%) | 695 (0.90%) | 2,849 (0.17%) | 16,695 (0.17%) |

**Supplemental Table 2. Performance comparison of KGIPA with other state-of-the-art methods based on five-fold cross-validation.**

| Methods | Evaluation Metrics | | | | |
|---|---|---|---|---|---|
| | **MCC ↑** | **ACC ↑** | **F1 ↑** | **AUROC ↑** | **AUPR ↑** |
| **Protein Binding Residue Identification** | | | | | |
| MONN | $0.490_{\pm 0.008}$ | $0.964_{\pm 0.001}$ | $0.509_{\pm 0.008}$ | $0.887_{\pm 0.003}$ | $0.468_{\pm 0.008}$ |
| IIDL-PepPI | $0.427_{\pm 0.010}$ | $0.967_{\pm 0.001}$ | $0.419_{\pm 0.014}$ | $0.881_{\pm 0.003}$ | $0.431_{\pm 0.010}$ |
| **KGIPA** | $\mathbf{0.552_{\pm 0.006}}$ | $\mathbf{0.968_{\pm 0.001}}$ | $\mathbf{0.569_{\pm 0.005}}$ | $\mathbf{0.941_{\pm 0.002}}$ | $\mathbf{0.577_{\pm 0.006}}$ |
| **Peptide Binding Residue Identification** | | | | | |
| CAMP | $0.357_{\pm 0.011}$ | $0.734_{\pm 0.005}$ | $0.545_{\pm 0.047}$ | $0.759_{\pm 0.006}$ | $0.607_{\pm 0.009}$ |
| MONN | $0.326_{\pm 0.013}$ | $0.653_{\pm 0.017}$ | $0.580_{\pm 0.007}$ | $0.741_{\pm 0.006}$ | $0.596_{\pm 0.011}$ |
| IIDL-PepPI | $0.380_{\pm 0.004}$ | $0.734_{\pm 0.002}$ | $0.569_{\pm 0.007}$ | $0.765_{\pm 0.003}$ | $0.632_{\pm 0.006}$ |
| **KGIPA** | $\mathbf{0.429_{\pm 0.005}}$ | $\mathbf{0.738_{\pm 0.002}}$ | $\mathbf{0.632_{\pm 0.003}}$ | $\mathbf{0.804_{\pm 0.003}}$ | $\mathbf{0.694_{\pm 0.003}}$ |

The results are presented as mean ± standard deviation. The best result for each metric is marked in bold.

**Supplemental Table 3. Performance comparison of KGIPA with protein language models for protein- and peptide-binding residue recognition based on independent test dataset Test167.**

| Methods | Evaluation Metrics | | | | |
|---|---|---|---|---|---|
| | MCC ↑ | ACC ↑ | F1 ↑ | AUROC ↑ | AUPR ↑ |
| **Protein Binding Residue Identification** | | | | | |
| TAPE | $0.226_{\pm 0.004}$ | $0.957_{\pm 0.003}$ | $0.247_{\pm 0.002}$ | $0.783_{\pm 0.001}$ | $0.194_{\pm 0.003}$ |
| ProtBert | $0.252_{\pm 0.002}$ | $0.952_{\pm 0.001}$ | $0.276_{\pm 0.001}$ | $0.787_{\pm 0.001}$ | $0.212_{\pm 0.001}$ |
| ESM 2-650M | $0.361_{\pm 0.003}$ | $0.963_{\pm 0.001}$ | $0.380_{\pm 0.003}$ | $0.862_{\pm 0.001}$ | $0.337_{\pm 0.002}$ |
| ESM C-600M | $0.353_{\pm 0.006}$ | $0.963_{\pm 0.002}$ | $0.372_{\pm 0.004}$ | $0.856_{\pm 0.001}$ | $0.329_{\pm 0.000}$ |
| ProtT5-XL-U50 | $0.372_{\pm 0.004}$ | $0.964_{\pm 0.002}$ | $0.390_{\pm 0.005}$ | $0.864_{\pm 0.002}$ | $0.344_{\pm 0.001}$ |
| **KGIPA** | $\mathbf{0.533_{\pm 0.007}}$ | $\mathbf{0.970_{\pm 0.001}}$ | $\mathbf{0.547_{\pm 0.007}}$ | $\mathbf{0.937_{\pm 0.005}}$ | $\mathbf{0.556_{\pm 0.004}}$ |
| **Peptide Binding Residue Identification** | | | | | |
| TAPE | $0.295_{\pm 0.006}$ | $0.633_{\pm 0.020}$ | $0.539_{\pm 0.002}$ | $0.720_{\pm 0.002}$ | $0.489_{\pm 0.002}$ |
| ProtBert | $0.237_{\pm 0.005}$ | $0.612_{\pm 0.016}$ | $0.504_{\pm 0.001}$ | $0.668_{\pm 0.001}$ | $0.441_{\pm 0.001}$ |
| ESM 2-650M | $0.337_{\pm 0.001}$ | $0.670_{\pm 0.006}$ | $0.561_{\pm 0.001}$ | $0.730_{\pm 0.001}$ | $0.498_{\pm 0.001}$ |
| ESM C-600M | $0.329_{\pm 0.005}$ | $0.659_{\pm 0.007}$ | $0.558_{\pm 0.003}$ | $0.731_{\pm 0.001}$ | $0.503_{\pm 0.002}$ |
| ProtT5-XL-U50 | $0.346_{\pm 0.002}$ | $0.693_{\pm 0.007}$ | $0.562_{\pm 0.002}$ | $0.737_{\pm 0.001}$ | $0.511_{\pm 0.003}$ |
| **KGIPA** | $\mathbf{0.473_{\pm 0.021}}$ | $\mathbf{0.750_{\pm 0.017}}$ | $\mathbf{0.643_{\pm 0.013}}$ | $\mathbf{0.845_{\pm 0.010}}$ | $\mathbf{0.709_{\pm 0.015}}$ |

The results are presented as mean ± standard deviation. The best result for each metric is marked in bold. ESM Cambrian (ESM C) is a parallel model family of ESM 3 generative models.

**Supplemental Table 4. Performance comparison of KGIPA with protein language models for protein- and peptide-binding residue recognition based on independent test dataset Test251&LEADS-PEP.**

| Methods | Evaluation Metrics | | | | |
|---|---|---|---|---|---|
| | **MCC ↑** | **ACC ↑** | **F1 ↑** | **AUROC ↑** | **AUPR ↑** |
| **Protein Binding Residue Identification** | | | | | |
| TAPE | $0.162_{\pm 0.002}$ | $0.907_{\pm 0.010}$ | $0.206_{\pm 0.002}$ | $0.720_{\pm 0.001}$ | $0.142_{\pm 0.001}$ |
| ProtBert | $0.165_{\pm 0.002}$ | $0.895_{\pm 0.006}$ | $0.208_{\pm 0.001}$ | $0.728_{\pm 0.001}$ | $0.139_{\pm 0.000}$ |
| ESM2-650M | $0.297_{\pm 0.003}$ | $0.926_{\pm 0.004}$ | $0.332_{\pm 0.003}$ | $0.823_{\pm 0.000}$ | $0.266_{\pm 0.001}$ |
| ESM C-600M | $0.280_{\pm 0.001}$ | $0.925_{\pm 0.004}$ | $0.316_{\pm 0.001}$ | $0.819_{\pm 0.001}$ | $0.245_{\pm 0.001}$ |
| ProtT5-XL-U50 | $0.320_{\pm 0.003}$ | $0.925_{\pm 0.002}$ | $0.352_{\pm 0.002}$ | $0.837_{\pm 0.001}$ | $0.295_{\pm 0.001}$ |
| **KGIPA** | $\mathbf{0.540_{\pm 0.006}}$ | $\mathbf{0.958_{\pm 0.002}}$ | $\mathbf{0.562_{\pm 0.005}}$ | $\mathbf{0.924_{\pm 0.002}}$ | $\mathbf{0.584_{\pm 0.010}}$ |
| **Peptide Binding Residue Identification** | | | | | |
| TAPE | $0.146_{\pm 0.015}$ | $0.546_{\pm 0.009}$ | $0.682_{\pm 0.001}$ | $0.659_{\pm 0.002}$ | $0.650_{\pm 0.001}$ |
| ProtBert | $0.015_{\pm 0.008}$ | $0.507_{\pm 0.001}$ | $0.673_{\pm 0.000}$ | $0.615_{\pm 0.001}$ | $0.620_{\pm 0.001}$ |
| ESM 2-650M | $0.282_{\pm 0.019}$ | $0.636_{\pm 0.013}$ | $0.689_{\pm 0.000}$ | $0.695_{\pm 0.001}$ | $0.690_{\pm 0.001}$ |
| ESM C-600M | $0.208_{\pm 0.005}$ | $0.587_{\pm 0.003}$ | $0.685_{\pm 0.001}$ | $0.679_{\pm 0.001}$ | $0.675_{\pm 0.002}$ |
| ProtT5-XL-U50 | $0.295_{\pm 0.004}$ | $0.638_{\pm 0.004}$ | $0.701_{\pm 0.002}$ | $0.712_{\pm 0.001}$ | $0.710_{\pm 0.001}$ |
| **KGIPA** | $\mathbf{0.339_{\pm 0.013}}$ | $\mathbf{0.652_{\pm 0.008}}$ | $\mathbf{0.722_{\pm 0.003}}$ | $\mathbf{0.761_{\pm 0.001}}$ | $\mathbf{0.773_{\pm 0.004}}$ |

The results are presented as mean ± standard deviation. The best result for each metric is marked in bold. ESM Cambrian (ESM C) is a parallel model family of ESM 3 generative models.

**Supplemental Table 5. Performance comparison of KGIPA with SOTA baselines and protein language models for residue-level interaction recognition based on independent test dataset Test1440.**

| Methods | Evaluation Metrics | | | | |
|---|---|---|---|---|---|
| | **MCC ↑** | **ACC ↑** | **F1 ↑** | **AUROC ↑** | **AUPR ↑** |
| **Protein Binding Residue Identification** | | | | | |
| RoseTTAFold | 0.217 | 0.922 | 0.233 | 0.660 | 0.284 |
| PepNN | 0.218 | 0.941 | 0.243 | 0.814 | 0.160 |
| TAPE | $0.207_{\pm 0.003}$ | $0.954_{\pm 0.001}$ | $0.230_{\pm 0.002}$ | $0.782_{\pm 0.001}$ | $0.174_{\pm 0.001}$ |
| ProtBert | $0.254_{\pm 0.002}$ | $0.955_{\pm 0.002}$ | $0.277_{\pm 0.002}$ | $0.786_{\pm 0.001}$ | $0.209_{\pm 0.001}$ |
| ESM2-650M | $0.353_{\pm 0.001}$ | $0.962_{\pm 0.002}$ | $0.373_{\pm 0.001}$ | $0.860_{\pm 0.001}$ | $0.320_{\pm 0.001}$ |
| ESM C-600M | $0.345_{\pm 0.001}$ | $0.956_{\pm 0.001}$ | $0.366_{\pm 0.001}$ | $0.853_{\pm 0.001}$ | $0.309_{\pm 0.001}$ |
| ProtT5-XL-U50 | $0.356_{\pm 0.001}$ | $0.962_{\pm 0.001}$ | $0.375_{\pm 0.001}$ | $0.871_{\pm 0.001}$ | $0.324_{\pm 0.001}$ |
| MONN | $0.213_{\pm 0.015}$ | $0.970_{\pm 0.000}$ | $0.107_{\pm 0.013}$ | $0.843_{\pm 0.004}$ | $0.320_{\pm 0.015}$ |
| IIDL-PepPI | $0.273_{\pm 0.009}$ | $0.970_{\pm 0.001}$ | $0.218_{\pm 0.007}$ | $0.841_{\pm 0.009}$ | $0.326_{\pm 0.013}$ |
| AlphaFold3 | 0.406 | 0.951 | 0.415 | 0.760 | 0.451 |
| **KGIPA** | $\mathbf{0.527_{\pm 0.007}}$ | $\mathbf{0.972_{\pm 0.001}}$ | $\mathbf{0.542_{\pm 0.007}}$ | $\mathbf{0.933_{\pm 0.004}}$ | $\mathbf{0.532_{\pm 0.007}}$ |
| **Peptide Binding Residue Identification** | | | | | |
| RoseTTAFold | 0.103 | 0.558 | 0.403 | 0.558 | 0.496 |
| TAPE | $0.338_{\pm 0.004}$ | $0.703_{\pm 0.006}$ | $0.538_{\pm 0.002}$ | $0.735_{\pm 0.002}$ | $0.509_{\pm 0.001}$ |
| ProtBert | $0.269_{\pm 0.002}$ | $0.644_{\pm 0.008}$ | $0.500_{\pm 0.001}$ | $0.701_{\pm 0.001}$ | $0.462_{\pm 0.002}$ |
| ESM 2-650M | $0.339_{\pm 0.001}$ | $0.708_{\pm 0.006}$ | $0.537_{\pm 0.001}$ | $0.740_{\pm 0.001}$ | $0.515_{\pm 0.001}$ |
| ESM C-600M | $0.334_{\pm 0.001}$ | $0.703_{\pm 0.004}$ | $0.534_{\pm 0.002}$ | $0.741_{\pm 0.001}$ | $0.510_{\pm 0.001}$ |
| ProtT5-XL-U50 | $0.342_{\pm 0.002}$ | $0.709_{\pm 0.001}$ | $0.539_{\pm 0.002}$ | $0.745_{\pm 0.001}$ | $0.520_{\pm 0.001}$ |
| CAMP | $0.362_{\pm 0.006}$ | $0.774_{\pm 0.001}$ | $0.481_{\pm 0.010}$ | $0.778_{\pm 0.002}$ | $0.565_{\pm 0.004}$ |
| MONN | $0.228_{\pm 0.010}$ | $0.754_{\pm 0.001}$ | $0.185_{\pm 0.029}$ | $0.762_{\pm 0.005}$ | $0.564_{\pm 0.003}$ |
| IIDL-PepPI | $0.368_{\pm 0.006}$ | $0.766_{\pm 0.005}$ | $0.517_{\pm 0.008}$ | $0.769_{\pm 0.006}$ | $0.572_{\pm 0.013}$ |
| AlphaFold3 | 0.301 | 0.680 | 0.515 | 0.666 | 0.583 |
| **KGIPA** | $\mathbf{0.473_{\pm 0.006}}$ | $\mathbf{0.780_{\pm 0.004}}$ | $\mathbf{0.623_{\pm 0.004}}$ | $\mathbf{0.831_{\pm 0.005}}$ | $\mathbf{0.675_{\pm 0.006}}$ |
| **Pairwise Interaction Residue Identification** | | | | | |
| RoseTTAFold | 0.094 | 0.990 | 0.072 | 0.606 | 0.133 |
| MONN | $0.138_{\pm 0.009}$ | $0.998_{\pm 0.000}$ | $0.063_{\pm 0.007}$ | $0.879_{\pm 0.009}$ | $0.096_{\pm 0.006}$ |
| AlphaFold3 | 0.261 | 0.995 | 0.233 | 0.716 | **0.298** |
| **KGIPA** | $\mathbf{0.324_{\pm 0.005}}$ | $\mathbf{0.998_{\pm 0.000}}$ | $\mathbf{0.325_{\pm 0.004}}$ | $\mathbf{0.949_{\pm 0.005}}$ | $0.256_{\pm 0.004}$ |

The results are presented as mean ± standard deviation. The best result for each metric is marked in bold. ESM Cambrian (ESM C) is a parallel model family of ESM 3 generative models.

**Supplemental Table 6. Comparison of computational cost and predictive performance between KGIPA and baseline methods.**

| Method | Training (s/epoch) ↓ | Inference (s/pair) ↓ | Score ↑ |
|---|---|---|---|
| **Not using structural features** | | | |
| ESM-2 | 45.40 | 1.11 | 0.418 |
| ESM-3 | 40.03 | 1.21 | 0.416 |
| IIDL-PepPI | 146.20 | 7.72 | 0.534 |
| KGIPA-Fast | 261.33 | 11.12 | 0.607 |
| **Using structural features (predicted or experimentally determined)** | | | |
| RFAA | / | 372.75 | 0.369 |
| AlphaFold3 | / | 109.69 | 0.541 |
| KGIPA | 325.45 | 111.38 | 0.633 |

Inference time includes the feature extraction process for each method. Score represents the mean AUPR for predicted peptide and protein binding residues.

**Supplemental Table 7. Feature ablation analysis of the KGIPA model.**

| Methods | Evaluation Metrics | | | | |
|---|---|---|---|---|---|
| | MCC ↑ | ACC ↑ | F1 ↑ | AUROC ↑ | AUPR ↑ |
| **Protein Binding Residue Identification** | | | | | |
| **KGIPA** | **0.552±0.006** | **0.968±0.001** | **0.569±0.005** | **0.941±0.002** | **0.577±0.006** |
| - trRosetta | 0.541±0.006 | 0.966±0.001 | 0.559±0.006 | 0.940±0.002 | 0.563±0.012 |
| - ProtT5 | 0.539±0.003 | 0.965±0.001 | 0.556±0.002 | 0.939±0.001 | 0.556±0.002 |
| - PSI-BLAST | 0.545±0.003 | 0.967±0.001 | 0.562±0.003 | 0.940±0.002 | 0.568±0.006 |
| - IUPred2A | 0.540±0.009 | 0.966±0.001 | 0.558±0.008 | 0.939±0.002 | 0.559±0.013 |
| - SCRATCH | 0.543±0.004 | 0.966±0.001 | 0.560±0.004 | 0.940±0.003 | 0.564±0.011 |
| **Peptide Binding Residue Identification** | | | | | |
| **KGIPA** | **0.429±0.005** | **0.738±0.002** | 0.632±0.003 | **0.804±0.003** | **0.694±0.003** |
| - trRosetta | **0.429±0.005** | 0.725±0.006 | **0.634±0.003** | 0.803±0.003 | 0.687±0.005 |
| - ProtT5 | 0.402±0.007 | 0.704±0.008 | 0.620±0.006 | 0.789±0.001 | 0.670±0.005 |
| - PSI-BLAST | 0.426±0.005 | 0.725±0.006 | 0.631±0.003 | 0.802±0.002 | 0.686±0.003 |
| - IUPred2A | 0.425±0.010 | 0.725±0.012 | 0.629±0.005 | 0.801±0.003 | 0.685±0.009 |
| - SCRATCH | 0.425±0.008 | 0.720±0.005 | 0.631±0.005 | 0.803±0.004 | 0.689±0.004 |

The results are presented as mean ± standard deviation. The best result for each metric is marked in bold.

**Supplemental Table 8. Comparison of measured binding free energy changes ($\Delta\Delta G$) and KGIPA-predicted binding score changes ($\Delta Score$) for p53 phosphorylation mutants at Ser15 and Thr18 with different protein partners.**

| Interaction | Method | Ser15P | Thr18P |
|---|---|---|---|
| p53-MDM2 | Measured $\Delta\Delta G$ (kcal/mol) | - (0.00) | ↓ (1.11) |
| | Predicted $\Delta Score$ (%) | ↓ (1.05) | ↓ (1.09) |
| p53-p300 | Measured $\Delta\Delta G$(kcal/mol) | ↑ (-0.64) | ↑ (-1.15) |
| | Predicted $\Delta Score$ (%) | ↑ (-1.12) | ↑ (-1.38) |

Compared with the wild type, "↑" indicates that the mutation favors binding, "↓" indicates that the mutation disfavors binding, and "-" indicates little or no effect on binding.

**Supplemental Table 9. Detailed information on peptide-protein non-covalent interactions collected in this study.**

| Type | Protein Side | | Peptide Side | |
|---|---|---|---|---|
| | Number of positive samples | Percentage of positive samples | Number of positive samples | Percentage of positive samples |
| Hydrophobic Interaction | 41,015 | 1.782% | 28,231 | 17.551% |
| Hydrogen Bond | 41,270 | 1.793% | 32,721 | 20.343% |
| Water Bridge | 11,176 | 0.486% | 9,820 | 6.105% |
| Salt Bridge | 6,655 | 0.289% | 5,948 | 3.697% |
| π stacks | 816 | 0.035% | 783 | 0.487% |
| π-cation interactions | 978 | 0.042% | 867 | 0.539% |
| Halogen Bonds | 9 | 0.004‰ | 9 | 0.006% |
| Total | 84,539 | 3.674% | 52,288 | 32.508% |

**Supplemental Table 10. Performance comparison of KGIPA and IIDL-PepPI under identical feature inputs.**

| Methods | Evaluation Metrics | | | | |
|---|---|---|---|---|---|
| | **MCC ↑** | **ACC ↑** | **F1 ↑** | **AUROC ↑** | **AUPR ↑** |
| **Protein Binding Residue Identification** | | | | | |
| **KGIPA** | **0.552$_{\pm 0.006}$** | **0.968$_{\pm 0.001}$** | **0.569$_{\pm 0.005}$** | **0.941$_{\pm 0.002}$** | **0.577$_{\pm 0.006}$** |
| KGIPA Comparison | 0.537$_{\pm 0.007}$ | 0.966$_{\pm 0.001}$ | 0.555$_{\pm 0.007}$ | 0.938$_{\pm 0.003}$ | 0.552$_{\pm 0.011}$ |
| IIDL-PepPI | 0.427$_{\pm 0.010}$ | 0.967$_{\pm 0.001}$ | 0.419$_{\pm 0.014}$ | 0.881$_{\pm 0.003}$ | 0.431$_{\pm 0.010}$ |
| **Peptide Binding Residue Identification** | | | | | |
| **KGIPA** | **0.429$_{\pm 0.005}$** | **0.738$_{\pm 0.002}$** | 0.632$_{\pm 0.003}$ | **0.804$_{\pm 0.003}$** | **0.694$_{\pm 0.003}$** |
| KGIPA Comparison | 0.409$_{\pm 0.006}$ | 0.713$_{\pm 0.004}$ | 0.623$_{\pm 0.002}$ | 0.792$_{\pm 0.006}$ | 0.673$_{\pm 0.007}$ |
| IIDL-PepPI | 0.380$_{\pm 0.004}$ | 0.734$_{\pm 0.002}$ | 0.569$_{\pm 0.007}$ | 0.765$_{\pm 0.003}$ | 0.632$_{\pm 0.006}$ |

### Supplementary Notes

#### Supplemental Note 1: Generalization assessment across novel peptide, protein, and pairwise settings

To evaluate the model under more stringent scenarios, we further conducted generalization experiments based on Smith-Waterman (SW) local alignment for three cases: novel peptides, novel proteins, and novel peptide-protein pairs. SW uses dynamic programming for local alignment, providing a stricter assessment of sequence similarity at the residue level, which is particularly suitable for analyzing peptide-protein interactions at this granularity. Specifically, in the novel peptide scenario, we clustered peptide sequences using different SW thresholds, randomly partitioned them into three clusters of comparable size, and performed three-fold cross-validation to assess generalization performance on the peptide binding residue identification task. In the novel protein scenario, the same strategy was applied to protein sequences for evaluating protein binding residue identification. Furthermore, in the novel peptide-protein pair scenario, we combined peptide clusters and protein clusters in a pairwise manner to form nine data clusters and carried out nine-fold cross-validation to evaluate the generalization ability of MONN and KGIPA on the pairwise binding residue identification task. The results show that as the clustering becomes stricter (i.e., higher SW thresholds), the generalization performance of all methods decreases, yet KGIPA consistently maintains superior performance across all three scenarios (Figure S1). This demonstrates its ability to effectively capture peptide-protein interaction patterns and generalize well to unseen data. It is worth emphasizing that CAMP only supports peptide binding residue identification, while MONN and KGIPA are the only methods capable of predicting pairwise residue interactions. Moreover, since structure prediction methods such as RoseTTAFold and AlphaFold3 do not allow model retraining, they were not included in these experiments.

#### Supplemental Note 2: Computational efficiency analysis of KGIPA and baseline methods

We evaluated KGIPA alongside several representative baseline methods to assess their computational efficiency and predictive performance. The baseline methods include models that leverage pretrained protein language model embeddings, such as ESM-2 and ESM-3 combined with fully connected layers, more advanced architectures such as IIDL-PepPI, and recent structure-prediction frameworks such as RoseTTAFold All-Atom (RFAA) and AlphaFold3. For each method, we measured the time consumption during training and inference and reported their predictive performance on the Test167 dataset, as shown in Table S6. All experiments were conducted on a workstation equipped with an Intel Platinum 8378A CPU (128 logical cores) and eight NVIDIA A800-SXM4-40GB GPUs.

During the training phase, KGIPA exhibits a longer per-epoch runtime than baseline models. This is primarily because its contextualized interaction representation module needs to model multiple types of residue-level pairwise interaction graphs, inherently increasing computational complexity. Although methods such as ESM-2 or ESM-3 combined with fully connected layers train more rapidly, these methods do not account for the extensive training cost of large language models themselves. Moreover, since models such as RFAA and AlphaFold3 provide only pretrained models or online prediction services, their training time cannot be fairly evaluated.

In the inference phase, our analysis shows that the major computational overhead of KGIPA arises from molecular representation generation, particularly from the use of trRosetta to predict 3D structural features and SCRATCH to extract 2D structural descriptors. Consistent with this observation, methods that make use of structural features—whether experimentally determined or predicted—such as RFAA and AlphaFold3 generally require substantially more computational time than methods that do not utilize structural features, for example ESM-2 and ESM-3 combined with fully connected layers. Nevertheless, KGIPA achieves comparable inference efficiency to AlphaFold3 (≈111.38s vs. ≈109.69s) while delivering superior predictive performance.

To mitigate the relatively high computational cost of KGIPA during preprocessing, training, and inference, we developed a parallelized variant, KGIPA-Fast, by removing the 3D structural features predicted by trRosetta and the secondary structural features obtained from SCRATCH. Experimental results show that excluding structural information substantially reduces training and inference time, with only a minor decrease in prediction accuracy (Table S7). Researchers can thus select the full KGIPA model or the fast variant according to their application scenarios and computational needs. The KGIPA-Fast variant has also been integrated into our web server (http://bliulab.net/KGIPA), and for high-throughput analyses, users can deploy the standalone version locally to perform parallel processing or model retraining (https://github.com/ShutaoChen97/KGIPA).

#### Supplemental Note 3: Supplementary insights into key residue identification

In our study, we conducted two independent sets of in silico alanine scanning analyses to evaluate the impact of mutating predicted binding residues. While the adjusted $R^2$ values (e.g., 0.827) may appear moderate, the Pearson correlation coefficients (e.g., 0.918) are notably higher, indicating that KGIPA effectively captures the trend of binding affinity changes rather than absolute $\Delta\Delta G$ values (Figure S3). This is particularly meaningful considering that our predictions were not fitted to any specific experimental alanine scanning dataset. Instead, they simulate general mutation effects across diverse peptide-protein complexes. This result suggests that KGIPA can reliably identify residues whose substitution tends to have greater or lesser functional impact, which is especially helpful for guiding the identification of key residues in biological experiments. In this context, the emphasis is on prioritizing mutation-sensitive sites, rather than reproducing precise energy changes, a strategy that aligns with early-stage experimental design.

In addition to the two alanine scanning experiments described above, we extended our investigation to a systematic analysis of cases where the same peptide engages with different protein partners, thereby providing further validation of the context sensitivity captured by KGIPA. Specifically, we selected the p53 transcriptional activation domain 1 (TAD1, 13-29 aa) as the study subject. This region, located at the N-terminus of p53, represents a core functional fragment known to interact with multiple regulatory proteins and plays a critical role in cell cycle control and stress responses. As this peptide often exhibits distinct binding modes and functional outcomes depending on its partners, it provides an ideal case for analysis. In terms of protein partners, we focused on two structurally and functionally divergent examples:

1) MDM2, the classical inhibitor of p53, which binds to TAD1 and mediates its ubiquitination and degradation, thereby serving as a key negative regulator of p53 activity.

2) p300 (Taz2 domain), an important transcriptional coactivator, which binds to TAD1 to promote transcriptional activation and enhance downstream gene expression.

Notably, these two proteins exhibit differences at the peptide-side binding interface when interacting with p53 TAD1. This experimental design therefore allows a more targeted comparison of the same peptide under different functional contexts, highlighting KGIPA's ability to capture interaction context dependency.

We first performed a residue-by-residue alanine scanning on the reported key binding region of this peptide (18-28 aa). This segment was chosen because it lies within TAD1 and has been demonstrated to be the most critical fragment for interactions with multiple binding partners, making it highly representative. Subsequently, we compared the changes in predicted binding probabilities ($\Delta Score$) between the wild type and each point mutant when interacting with MDM2 and p300 using KGIPA. Specifically, $\Delta Score$ is defined as:

$$\Delta Score = 1 + \frac{Score_{WT} - Score_{Ala}}{Score_{WT}} \tag{S1}$$

where $Score_{WT}$ denotes the predicted binding probability of the wild-type peptide, and $Score_{Ala}$ represents that of the alanine mutant. Accordingly, $\Delta Score > 1$ indicates that alanine substitution is unfavorable for binding (with larger values suggesting a stronger negative effect), whereas $\Delta Score < 1$ suggests that the mutation is, to some extent, favorable for binding. The results (Figure S4) show that even for the same peptide, KGIPA can capture binding interface variations arising from differences in protein-side environments, thereby demonstrating its context-dependent pragmatic effect.

Furthermore, we evaluated whether KGIPA could identify the key residues involved in the p53-MDM2 and p53-p300 interactions. It is worth noting that the identification of critical residues traditionally relies on wet-lab techniques such as alanine scanning, which are both time-consuming and costly. Therefore, the ability to accurately identify key residues through computational approaches is of considerable importance. To this end, we adopted a dry-wet combined strategy to analyze the residues predicted by KGIPA. First, we compared the changes in binding probability predicted by KGIPA ($\Delta Score$) with the $\Delta\Delta G$ values obtained from Rosetta alanine scanning filters The results demonstrated that upon residue-by-residue alanine substitution of p53, the trends in KGIPA's predicted binding probability changes were highly consistent with Rosetta's ΔΔG analysis ($p$-value = $2.697 \times 10^{-5}$ for the p53-MDM2 complex and $p$-value = $6.174 \times 10^{-4}$ for the p53-p300 complex), indicating that the model effectively distinguishes context-dependent key residues for the same peptide across different receptors. We further validated KGIPA's predictions against experimental measurements. In the p53-MDM2 interaction, experiments have shown that Phe19 ($\Delta\Delta G$ = 10.29 kcal/mol), Trp23 ($\Delta\Delta G$ = 6.81 kcal/mol), and Leu26 ($\Delta\Delta G$ = 2.39 kcal/mol) are the core residues driving binding, all of which were accurately reflected in KGIPA's predictions (Figure S5A). In contrast, for the p53-p300 interaction, the contributions of Trp23 and Leu26 were attenuated, whereas Leu22 ($\Delta\Delta G$ = 2.45 kcal/mol) and Leu25 ($\Delta\Delta G$ = 1.69 kcal/mol) emerged as the dominant hydrophobic binding residues, again consistent with KGIPA's predictions (Figure S5B). Together, these results demonstrate that KGIPA not only captures the environmental dependency of peptide-protein binding but also accurately identifies key residues under different receptor contexts, thereby validating its effectiveness in modeling biologically relevant context dependence.

Furthermore, to validate KGIPA's ability to capture context sensitivity in broader scenarios, we examined the effects of site-specific phosphorylation on the interactions of p53 with MDM2 and p300. Previous studies have shown that a key feature of the p53-p300 interaction is that its binding affinity can be markedly enhanced by phosphorylation at Ser15 and Thr18, whereas the p53-MDM2 interaction is largely unaffected. Based on this experimental evidence, we introduced phosphorylation at the corresponding sites to assess whether KGIPA could capture this effect. As shown in Table S8, the predictions revealed that phosphorylation at Ser15 and Thr18 strengthens the binding affinity in the p53-p300 interaction ($\Delta Score < 1$ indicates a mutation favorable for binding), while the same modifications exert weaker effects on the p53-MDM2 interaction and do not confer substantial enhancement. Moreover, when comparing KGIPA's predictions with experimentally measured $\Delta\Delta G$ values ($\Delta\Delta G < 0$ indicates a mutation favorable for binding, whereas $\Delta\Delta G > 0$ indicates unfavorable effects), the two are largely consistent in trend. It is worth noting that in the case of Ser15 phosphorylation in the p53-MDM2 interaction, experimental measurements reported no change in binding affinity, whereas KGIPA predicted a slight decrease. This discrepancy may stem from experimental conditions or limitations in measurement precision. Nevertheless, the predicted decrease was minimal (with $\Delta Score$ approaching the threshold of 1 used to distinguish favorable from unfavorable effects) and is practically negligible, thus not altering the overall conclusion.

#### Supplemental Note 4: Comparison between KGIPA and IIDL-PepPI

The differences between IIDL-PepPI and KGIPA can be summarized in three major aspects: conceptual framing, problem definition, and methodological implementation. At the conceptual level, KGIPA deepens and systematizes the notion of pragmatic analysis first proposed by IIDL-PepPI. At the problem level, KGIPA extends the predictive scope from binary interactions and binding residues to residue-residue interaction maps and diverse non-covalent mechanisms. At the implementation level, KGIPA introduces bilinear modeling and knowledge-guided modules that improve biological relevance. The detailed distinctions are outlined below.

Conceptual level: IIDL-PepPI first introduced the idea of pragmatic analysis, emphasizing that peptide-protein interaction predictions should consider the binding partner, i.e., the context dependency. Building on this concept, KGIPA further refines and systematizes pragmatic analysis by decomposing it into intra-linguistic context representation and extra-linguistic context representation. Intra-linguistic context focuses on integrating multimodal features within a single molecule, while extra-linguistic context models feature interactions across multiple molecules. This design aligns with the biological rationale of "analyzing the monomer before the environment" and is analogous to the natural language processing principle of "analyzing the sentence before the full text." Therefore, conceptually, KGIPA represents a deepening and expansion of IIDL-PepPI.

Task level: IIDL-PepPI primarily aims to predict binary peptide-protein interactions and identify binding residues, i.e., answering "whether binding occurs" and "which residues are important." KGIPA extends this framework by predicting residue-residue interaction maps, capturing pairwise relationships between peptide and protein residues and dissecting multiple types of non-covalent interaction mechanisms. This extension from single-point predictions to edge-level modeling not only enhances interpretability but also provides more precise guidance for related biological experiments. In terms of application, IIDL-PepPI is suitable for large-scale candidate peptide screening, whereas KGIPA offers advantages in functional interpretation, drug design, and analyses of modifications or mutational effects, highlighting its expanded task capabilities and practical utility.

Implementation level: IIDL-PepPI mainly relies on a bidirectional attention mechanism for feature integration, yet its outputs remain limited to either peptide-side or protein-side predictions and lack direct interpretability of residue-residue interactions. KGIPA incorporates a bilinear network to explicitly model residue-residue interactions, establishing one-to-one correspondences between peptide and protein residues, which substantially aligns predictions more closely with biological processes. Additionally, KGIPA introduces a knowledge-guided module that simultaneously models multiple non-covalent interactions, including hydrogen bonds, hydrophobic interactions, and electrostatic interactions, while capturing their cooperative or competitive relationships. This incorporation of biophysical priors further enhances the model's ability to capture the underlying principles of molecular interactions.

In summary, KGIPA advances beyond IIDL-PepPI in conceptual design, task formulation, and methodological implementation, providing a more comprehensive framework for modeling peptide-protein interactions.

To further demonstrate that the reported improvements of KGIPA stem from its novel components rather than differences in data processing or model tuning, we conducted additional comparative experiments on the same dataset. Specifically, we retrained the model using the KGIPA architecture with the same feature inputs as IIDL-PepPI (based on ProtBert and excluding trRosetta-predicted 3D structural information), thereby isolating the contribution of the new components. The results are summarized in Table S10. It is important to note that both IIDL-PepPI and KGIPA results reported in the manuscript were retrained on the same curated dataset and optimized via grid search to minimize potential biases from data processing or hyperparameter selection. The results indicate that even with identical feature inputs, KGIPA still achieves superior predictive performance compared to IIDL-PepPI. Furthermore, we performed paired t-tests on the five-fold cross-validation AUPR scores of KGIPA and IIDL-PepPI. The results show that KGIPA significantly outperforms IIDL-PepPI in both tasks: protein-binding residue prediction ($p$-value = $7.24\times10^{-6}$) and peptide-binding residue prediction ($p$-value = $2.64\times10^{-5}$). Similarly, when comparing the KGIPA variant model (KGIPA Comparison, i.e., KGIPA architecture with IIDL-PepPI feature inputs) with IIDL-PepPI, statistically significant performance differences were observed for both tasks: protein-binding residue prediction ($p$-value = $1.41\times10^{-4}$) and peptide-binding residue prediction ($p$-value = $7.13\times10^{-4}$). These results collectively indicate that the performance gains of KGIPA primarily arise from its novel components rather than external factors.
